# Nintedanib and Pirfenidone Affect Growth and Differentiation of Human Alveolar Type 2 Cells

**DOI:** 10.64898/2026.03.02.708135

**Authors:** Alexey Bazarov, Andrea Serra-Marques, Giulia Protti, Monica Yang, Ram Naikawadi, Gary Green, Seoyeon Lee, Jasleen Kukreja, Michael A. Matthay, Michael Wax, Xiaodan Cai, Rachel M. Wolters, Jason R. Rock, David Garfield, Paul J. Wolters

## Abstract

**Background:** Idiopathic pulmonary fibrosis (IPF) is a progressive fibrotic lung disease characterized by epithelial cell senescence. Pirfenidone and nintedanib are approved drugs for the treatment of IPF. They significantly slow disease progression, but their mechanisms of action, especially on alveolar type 2 (AT2) cells, are poorly understood. We addressed this question by evaluating colony formation and growth of human AT2 cells co-cultured with fibroblasts in organoid culture in the presence of pirfenidone and nintedanib. We further evaluated molecular changes induced by these drugs via single cell RNA-seq of treated organoids.

**Methods:** AT2 cells isolated from normal donor lungs or IPF patients were mixed with human fibroblasts in 3D culture and grown in the absence or presence of pirfenidone or nintedanib. After 14 days in culture, the organoids were quantified and cells extracted from Matrigel for single cell RNA-seq.

**Results:** AT2 cell organoids cultured in the presence of pirfenidone or nintedanib resulted in increased colony formation and, in the case of nintedanib, in larger colonies. We observed that untreated or pirfenidone treated AT2 cells lost surfactant protein C (SFTPC) expression and acquired an expression profile consistent with keratin (KRT)17^high^/KRT5^−^ basaloid cells, whereas a larger proportion of nintedanib treated cells retained SFTPC expression. In contrast, AT2 cells treated with TGFβ inhibitor exhibited intermediate (SFTPC^−^/KRT17^low^) gene expression profile.

**Conclusion:** These results suggest that nintedanib maintains an AT2-like expression state in culture and acts proximal to TGFβ.

**Conflict of Interest Statement:** PJW was supported by grants from Boehringer Ingelheim, Roche, Sanofi, Pliant and Arda Therapeutics and received personal fees from Boehringer Ingelheim and Sanofi. None of these companies had a role in the design or analysis of the study or in the writing of the manuscript. ASM, GP, JRR and DG are employees of Genetech. The other authors have no conflicts of interest to declare.

## INTRODUCTION

Idiopathic pulmonary fibrosis (IPF) is an invariably fatal age-related chronic lung disease characterized by progressive lung scarring that results in respiratory failure (1). The prevalence of IPF doubles every decade after the age of 50 and accounts for up to 1% of deaths in some countries (2). IPF is characterized by excessive deposition of pathologic extracellular matrix (ECM), which result in changes in lung architecture and reduced lung function (3). In fibrotic lung, alveolar fibroblasts differentiate into profibrotic fibroblasts which pathologically produce the excess matrix (4, 5).

A crucial stage in the initiation of IPF is failure of alveolar type 2 (AT2) cells, which serve as epithelial progenitor cells in the lung parenchyma (6). AT2 cells are capable of long-term self-renewal and differentiation to alveolar epithelial type I (AT1) cells, which are essential for gas exchange (7). Lungs of IPF patients exhibit significantly reduced number of AT2 cells (8) with shortened telomeres, even in patients without known telomere-related mutations (9, 10). Telomere dysfunction in AT2 cells leads to senescence, preventing their self-renewal and contributing to their depletion and fibroproliferation (11). In addition to AT2 cell failure, there is pathologic accumulation of a population of epithelial cells displaying an aberrant basal cell phenotype in the parenchyma of IPF lungs. The origin of these aberrant basal cells is believed to be due to transdifferentiation of AT2 cells, partially driven by TGFβ (12).

Two medications, pirfenidone and nintedanib, have been shown to slow IPF progression (13–19), however their mechanism of action is not fully understood. Although the binding partner of pirfenidone is unknown, it reportedly reduces TNFα and IL6 secretion by immune cells (20). Pirfenidone also reverses TGF-β1 mediated upregulation of Col-I, Col-III, α-SMA and fibronectin in fetal human lung fibroblasts and inhibits MRTF signaling in IPF fibroblasts (21, 22). Nintedanib is a tyrosine kinase inhibitor with more than 50 targets including PDGFR, FGFR, VEGFR1-3, SRK, LCK and Lyn (23). It affects MAPK, PI3K/AKT, JAK/STAT, TGF-β, VEGF and WNT/β-catenin signaling pathways (23). Which of these pathways is critical to slowing IPF progression remains uncertain. While AT2 cells play a central role in IPF progression, the effects of pirfenidone and nintedanib on them are largely unknown. Here we address this question by treating cultured AT2 cells derived from IPF patients and normal donors with these two agents.

## MATHERIALS AND METHODS

### Cells

MRC5 fibroblasts were obtained from UCSF Cell and Genome Engineering Core and maintained in DMEM media (*Gibco*, *Waltham, MA*), supplemented with 20% Fetal Bovine Serum, 1% Penicillin/Streptomycin and 1% GlutaMAX.

### Human lung tissues

The studies described in this paper were conducted according to the principles of the Declaration of Helsinki. Written informed consent was obtained from all subjects, and the study was approved by the University of California, San Francisco Institutional Review Board (IRB # 13-10738). Fibrotic lung tissues were obtained at the time of lung transplantation from patients with a diagnosis of idiopathic pulmonary fibrosis. Normal lung tissues were obtained from lungs rejected for transplantation by the Northern California Transplant Donor Network.

Clinicopathological details are provided in supplementary table S1.

### Three-dimensional organoid culture

Sorted CD45^−^EPCAM^+^LysoTracker red^high^ cells (5000) were mixed with MRC5 fibroblasts at passage 22-24 (50,000) in 45 μL of TEC plus media (Supplemental Methods). The cell suspension was mixed with 45 μL of Growth Factor Reduced Matrigel (*Gibco*, *Waltham, MA)*, plated on the upper trans-well chamber (*Costar, Corning, NY*) and allowed to solidify at 37°C for 10 min. 500 μL of TEC plus supplemented with 10μM of Y-27632 (*TOCRIS, Bristol, United Kingdom*), was added to the bottom trans-well chamber. After 48 h in culture, the media in the bottom chamber was changed to TEC plus without Y-27632.

Organoids were grown in presence of LysoTracker red (24) in order to detect AT2 cells in real time. Cultures were fed every 48 h and allowed to grow for 14 days when images of the wells were obtained using EVOS M5000 microscope (*Invitrogen, Carlsbad, CA*) and organoids counted with FIJI software.

### Differential expression and gene set enrichment analysis

To conduct differential expression analysis, we first pseudo-bulked our samples per cell type per sample, restricting our analyses to pseudo-bulks containing at least 50 cells. For each contrast of interest, we subset our pseudo-bulk dataset to the relevant samples and processed the results using limmaVoom as previously described (25). Gene set enrichment analyses were conducted using the Hallmark categories from MsigDB (26), Fast Gene Set Enrichment Analysis (fgsea) (27), and cameraPR (28).

### PROGENy pathway analysis

To summarize changes in gene expression at the pathway level, we utilized the PROGENy package (29). Briefly, we evaluated pseudo-bulk counts using a linear model contrast using the decoupleR package function run_mlm (30). The resulting weights of these models were then multiplied by the pathway weights to get pathway activity scores.

### CellChat analysis

Cross-talk between fibroblasts and epithelial cells was inferred using CellChat (31), which predicts intercellular communication patterns by analyzing the differential expression of ligands in fibroblast gene expression data. Fibroblasts were selected as senders and epithelial cells as receivers.

### Slingshot trajectory analysis

To more formally capture patterns of relatedness and connectivity across cells in our epithelial culture, we made use of the Slingshot package (32) to conduct a trajectory analysis in corrected PCA space. In our analysis, we used Slingshot to construct a minimum spanning tree using the fastMM-corrected principal components of our epithelial cell-only integration. For construction of this MST, we used defined cell-type annotations followed by the application of the principal curves method as implemented in Slingshot to obtaine smoothed trajectories and pseudo-times across all cells. The result was a single lineage rooted in AT2 cells.

### Cell cycle analyses

To identify changes in cell cycle as a function of cell type annotation and treatment, we leveraged Tricycle package (33), a transfer learning approach that annotates individual cells by their continuous position in the cell cycle based on high-resolution data from annotated cycling cells. We then binarized these continuous values into discrete cell cycle stages following the author’s guidelines with 0.5pi used for the start of S stage, pi as the start of G2M stage, and 1.75pi-0.25pi as the G1/G0 stage.

## RESULTS

### Pirfenidone and nintedanib increase AT2 cell colony forming efficiency

To investigate the effect of pirfenidone and nintedanib on human AT2 cells, we co-cultured AT2 cells obtained from normal donor or IPF lungs with MRC5 human lung fibroblasts (34) in the absence or presence of either pirfenidone or nintedanib. Both pirfenidone and nintedanib treatment resulted in a significant increase in the formation of lysotracker^+^ organoids derived from both IPF patients and normal donors (Fig 1a-b). Nintedanib-treated organoids appeared larger than controls and had more structured appearance with luminal space surrounded by epithelial cells (Fig 1a, 3c). Moreover, cell cycle analyses indicated that a larger portion of nintedanib-treated epithelial cells was in S-phase, compared to control or pirfenidone-treated cells suggesting that nintedanib increased proliferation of these cells (Fig S1)

**Fig. 1.**
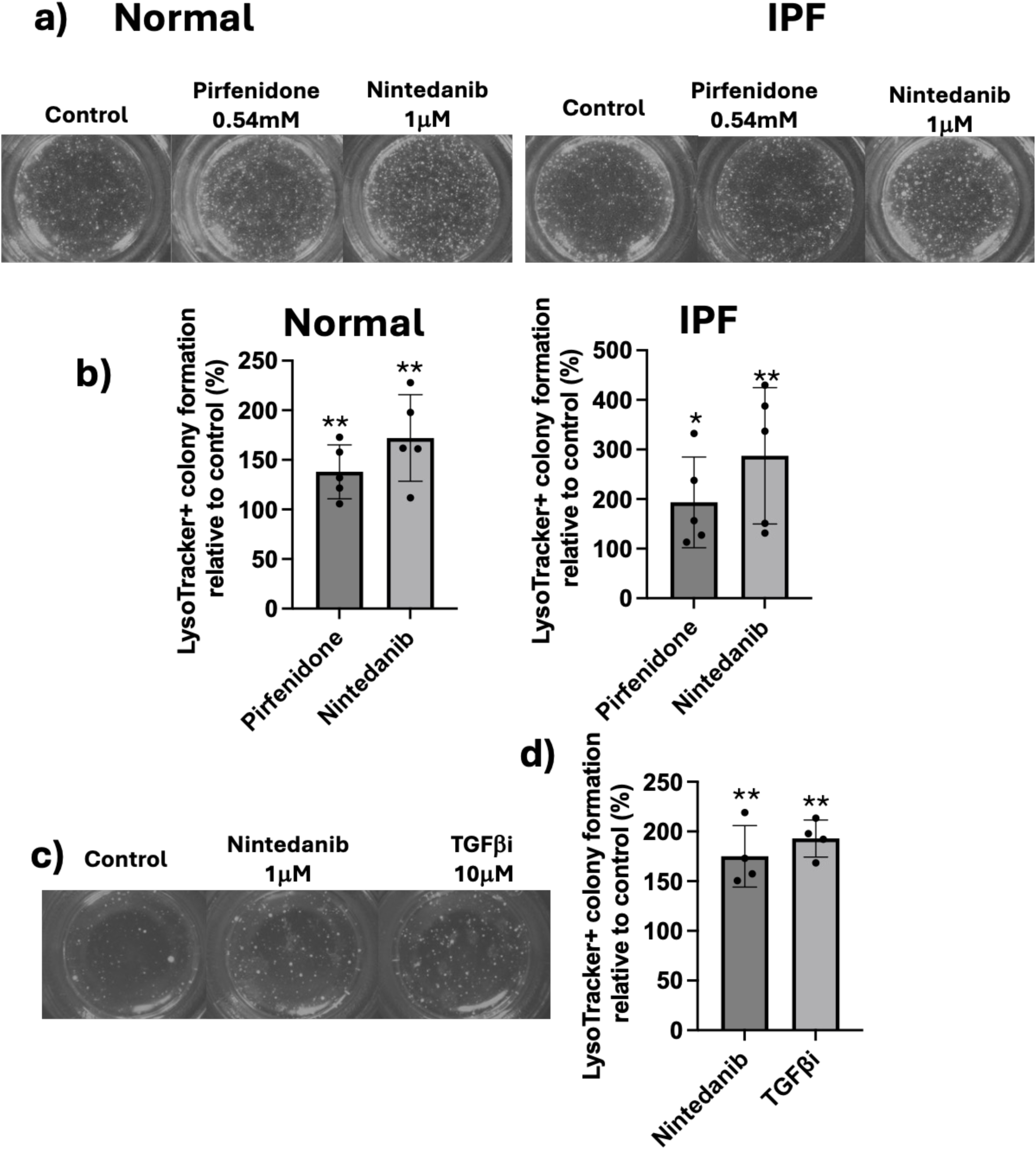
Pirfenidone, Nintedanib and TGF1 increase organoid formation in cultured AT2 cells. LysoTrackerhigh lung epithelial cells derived from either IPF patients or normal donors were co-cultured with MRC5 fibroblast for 2 weeks in continuous presence of Lysotracker red. **a)** Representative well images of lysotracker+ untreated organoids or treated with nintedanib or pirfenidone (1.25X). **b)** Quantification of lysotracker+ colony formation in pirfenidone and nintedanib treated cultures relative to controls. Plots show mean ± SD; n=5 normal and IPF samples. * p<0.05; ** p<0.01. **c)** Representative well images of lysotracker+ untreated organoids or treated with nintedanib or the TGFβ inhibitor SB-43152 (1.25X). **d**) Quantification of lysotracker+ colony formation in nintedanib or TGFβ inhibitor SB-43152 treated cultures relative to controls. Plots show mean ± SD of 3 replicas. ** p<0.01

TGFβ plays a critical role in IPF progression (35–38). We, therefore, tested whether inhibiting TGFβ would produce a similar effect to nintedanib and pirfenidone treatment. Culturing IPF-derived AT2 cells with the TGFβ inhibitor (TGFβi) SB43152 (39) resulted in a significant increase in lysotracker^+^ colony formation (Fig 1c-d).

### Nintedanib preserves AT2 cell markers in cultured LysoTracker-positive epithelial cells

To elucidate the mechanism of pirfenidone and nintedanib action, uncultured AT2 cells (day 0), and AT2 cell organoids co-cultured with fibroblasts in the absence or presence of pirfenidone, nintedanib, or TGFβi were analyzed by single-cell RNA sequencing (scRNA-seq) (Fig 2a). Because there was no significant difference in normal and IPF-derived cultured epithelial cells gene expression (Fig S2) and to achieve higher statistical power when evaluating the treatment groups, we combined the normal and IPF data sets. Epithelial cells were identified by EPCAM expression, while clusters lacking EPCAM and expressing PDGFRα were identified as fibroblasts (Fig. 2b-c). Uncultured AT2 cells clustered distinctly from cultured AT2 cells, as observed previously (40). Interestingly, while uncultured AT2 cells expressed high levels of SFTPC, only a subset of cultured epithelial cells did (Fig. 2d). Moreover, SFTPC expression was greatly reduced in control, pirfenidone– and TGFβi-treated cultured epithelial cells, while being relatively preserved in a subset of nintedanib-treated cells (Fig. 2e). This result was surprising, because pirfenidone, nintedanib, TGFβi-treated and controls cells all formed lysotracker-positive organoids (Fig. 1). As lysotracker binds to acidic lamellar bodies and is used as a marker of AT2 cells (41), the expression of surfactant proteins A1, A2, B and D (Fig. S3) was analyzed, finding that these genes were expressed in cultured epithelial cells regardless of treatment. Nintedanib treated cells also expressed higher levels of other AT2 markers (Fig. S4a.) and had a higher AT2 cell type gene score than other treatment conditions (Fig. 2g). In addition, nintedanib treatment resulted in the highest AT1 cell type gene score (Fig. S4c,f). Interestingly, TGFβi treated epithelial cells expressed higher levels of the Wnt-responsive AT2 cell markers TM4SF1 and FOXA1 compared to other treatments (Fig. S4e) (42).

**Fig. 2.**
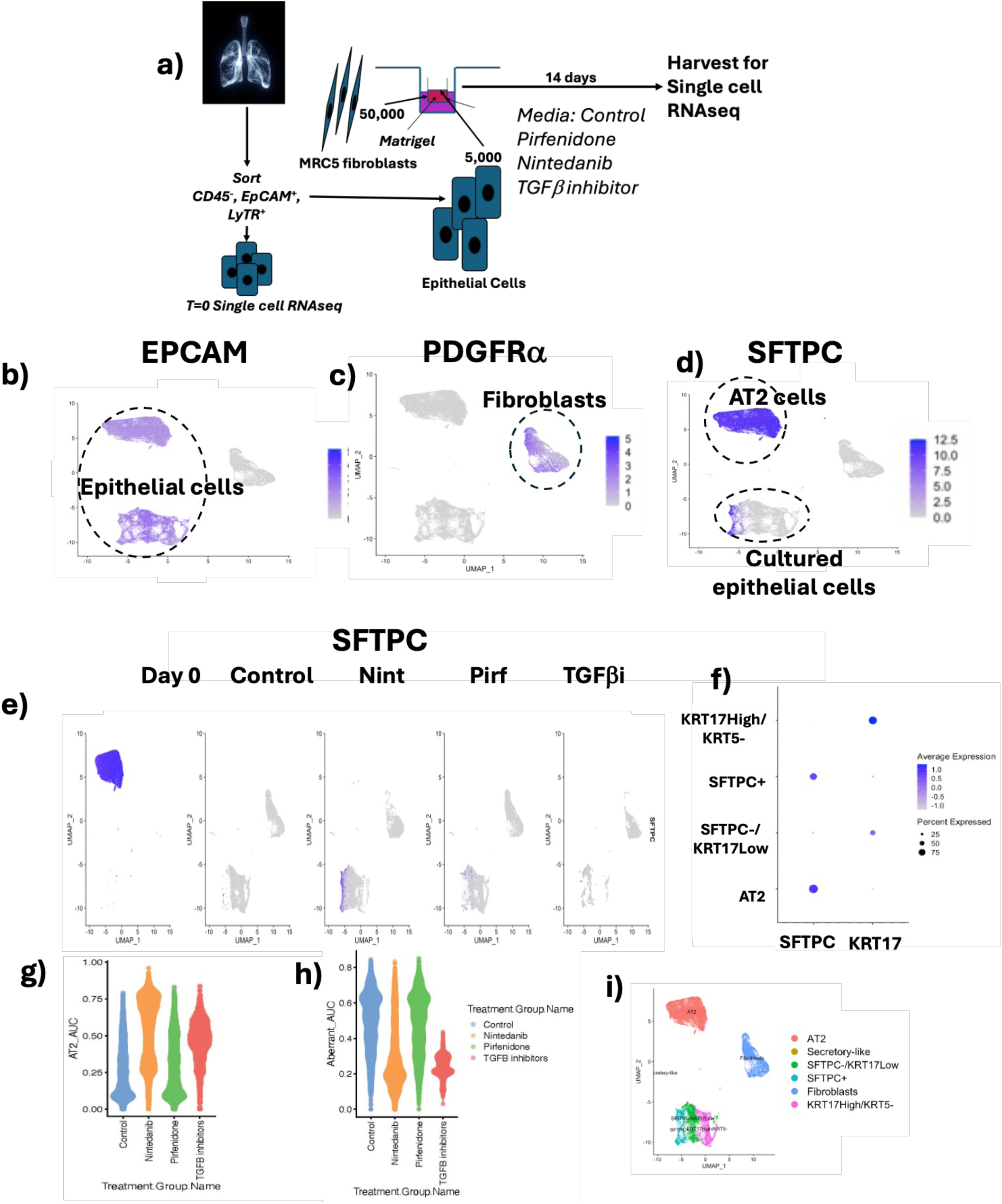
Clustering of cultured epithelial cells by scRNA-seq. CD45-/EPCAM+/LysoTrackerhigh cells were co-cultured with MRC5 fibroblasts for 14 days and analyzed by single cell RNA sequencing. **a)** Schematic of the scRNA-seq experiment (LyTR, LysoTracker Red). **b)** Feature plot for EPCAM **c)** Feature plot for SFTPC. **d)** Feature plot for PDGFRα **e)** SFTPC expression by treatment (Day 0: Freshly sorted cells prior to culture. **f)** Dotplot of SFTPC and KRT17 expression in epithelial cells. **g)** Area Under the Curve (AUC) score for AT2 markers. **h)** Area Under the Curve (AUC) score for aberrant basaloid markers. **i)** Clustering of epithelial cells and fibroblasts.

High levels of keratin 17 (KRT17) are associated with an abnormal population of basaloid epithelial cells, which are present in IPF but not in normal lungs (43). KRT17 was highly expressed in a subset of cultured epithelial cells that was distinct from SFTPC^+^ cultured cells (Fig. 2f). Cells expressing a high level of KRT17 were keratin 5 (KRT5) negative and also expressed TAGLN, LBH, MMP7, KRT7, KRT19, LAMC2, CDKN2a, TP63, FN1, CDH2, SOX4 and COL1A1 (Fig. S5), markers associated with aberrant basaloid cells (44). The aberrant cell type gene score was the highest in control and pirfenidone-treated cells (Fig. 2h, S4b). Senescence gene score was also the highest in these cells (Fig. S4 d,g). To investigate the relevance of the KRT17^high^/KRT5^−^ population in human disease, we constructed a gene signature composed of its top 30 differentially expressed genes relative to other epithelial cells. We then leveraged the SIGnature package, which allows to query a gene signature across more than 400 studies of various human diseases (45). This analysis revealed that the KRT17^high^/KRT5^−^ signature from our cell culture was enriched in pathological lung tissues, including those from patients with interstitial lung disease (ILD) and specifically IPF (Fig S6). The population located between SFTPC^+^ and KRT17^high^/KRT5^−^ cells expressed low levels of KRT17 and was labeled SFTPC^−^/KRT17^low^ (Fig. 2i).

To establish relationships between AT2 cells and the three populations of cultured epithelial cells that emerged in our data, we performed slingshot trajectory analysis. The results were consistent with a single trajectory leading from AT2 cells to KRT17^high^/KRT5^−^ cells through two intermediates SFTPC^+^ and SFTPC^−^/KRT17^low^ (Fig 3a). The distribution of cultured epithelial cells among these three groups varied with drug treatment. SFTPC^+^ cells were present almost exclusively in nintedanib-treated samples. TGFβi-treated organoids adopted transitional SFTPC^−^/KRT17^low^ phenotype, while control and pirfenidone-treated organoids yielded predominantly KRT17^high^/KRT5^−^ abnormal basaloid cells (Fig 3b-c). In agreement with single cell sequencing data, immunofluorescence analyses indicated that control organoids exhibited high levels of KRT17 staining, nintedanib-treated organoids high levels of SFTPC staining, while TGFβi treatment led to reductions of both SFTPC and KRT17 expression (Fig 3d).

**Fig. 3.**
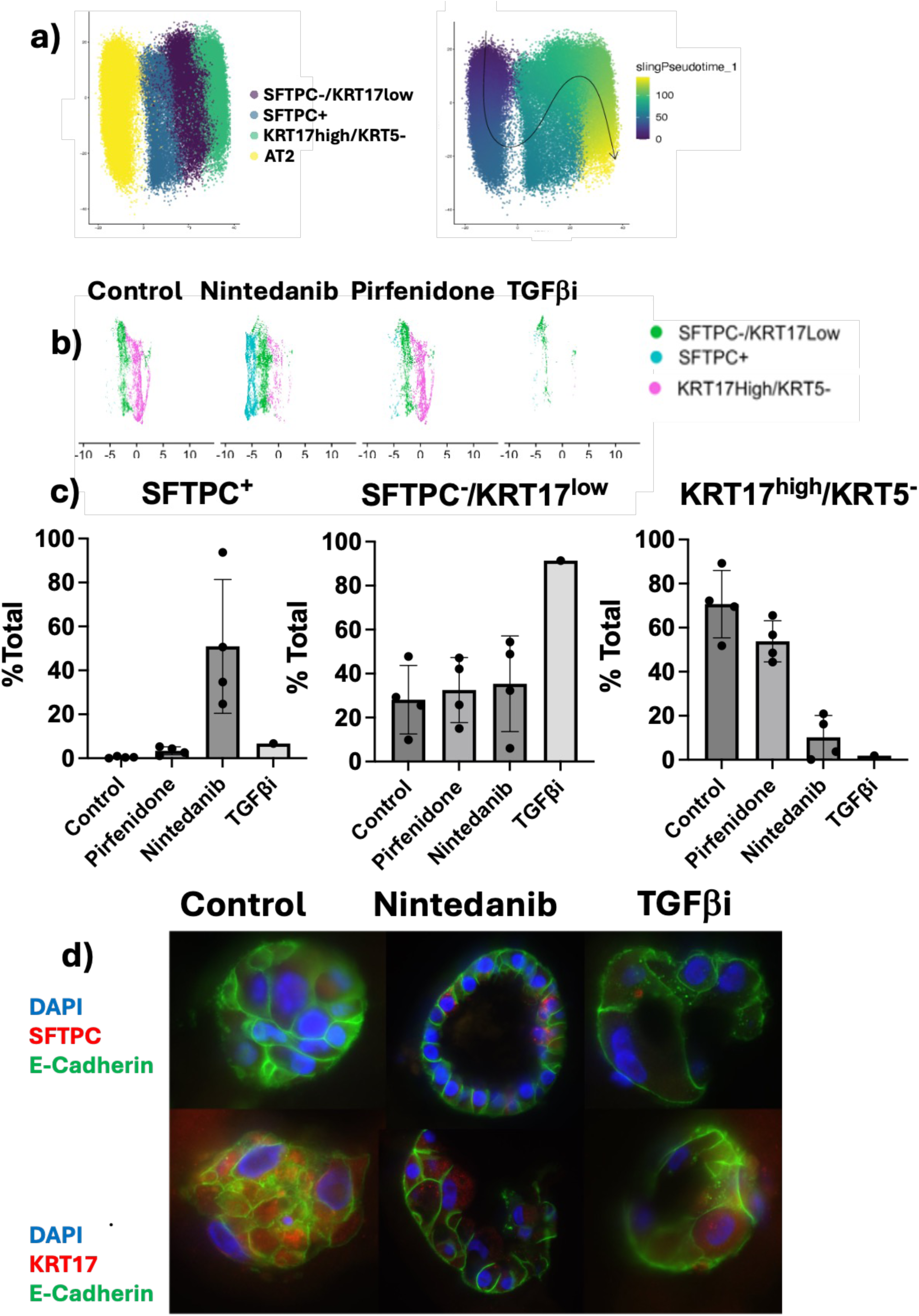
Nintedanib and TGFβ inhibitor arrest AT2 cell differentiation at distinct states. CD45-/EPCAM+/LysoTrackerhigh cells were co-cultured with MRC5 fibroblasts for 14 days and analyzed by single cell RNA sequencing. **a)** Slingshot trajectory analysis of epithelial cells **b)** Dimplot of cultured epithelial cells by conditions. **c)** Fraction of SFTPC^+^, SFTPC^−^/KRT17^low^ and KRT17^high^/KRT5^−^ presented as % total cultured epithelial cells. **d)** Representative IF images of organoids stained for SFTPC, KRT17 and E-Cadherin (40X magnification).

### Pirfenidone does not produce major changes in gene expression in co-cultured epithelial cells and fibroblasts

Pirfenidone significantly increased colony formation in normal and IPF-derived AT2 cell organoids (Fig 1). In addition, pirfenidone-treated epithelial cells slightly, but significantly, increased the population of SFTPC^+^ epithelial cells (3.36% Pirfenidone-treated, 0.46% Control, *p=0.011*) (Fig 3c). Despite these differences, there were no statistically significant changes in expression at a single gene level in pirfenidone-treated epithelial cells or fibroblasts compared to control (Fig S7a). Also, Milo analyses failed to reveal any significant differences between pirfenidone treated epithelial cells and controls (Fig S7b). One possible explanation is that pirfenidone acts through small changes in the expression of multiple genes, but the changes are sufficiently small that they do not reach statistical significance in these experiments. Consistent with this impression, Gene Set Enrichment Analysis (GSEA) revealed that pirfenidone caused significant pathway enrichment in both cultured fibroblasts and epithelial cells (Fig S8), indicating consistency in the sub significant changes in gene expression. EMT and p53 pathways were downregulated in pirfenidone-treated epithelial cells, while E2F targets and G2 checkpoint were upregulated. In pirfenidone-treated fibroblasts, fatty acid metabolism and MTOR signaling pathways were downregulated, as well as p53 and DNA repair pathways.

### Differential gene expression in nintedanib-treated cultured epithelial cells

Milo analyses reveal multiple significant changes in nintedanib-treated epithelial cells compared to controls (Fig. S9). The majority of SFTPC+ cells, which are the most transcriptionally similar to endogenous AT2 cells, came from nintedanib-treated cultures. However, despite a clear increase in the expression of AT2 markers relative to untreated controls, SFTPC+ cells showed numerous differences from pre-cultured AT2 cells. TGFβ signaling, TNFα via NFκB signaling, Wnt/β-catenin signaling as well as p53 pathway and UV response were downregulated in these cells (Fig. 4a). As compared to SFTPC^−^/KRT17^low^ epithelial cells, the SFTPC^+^ epithelial cells exhibit downregulation of EMT pathway, inflammatory response pathway, myogenesis, hypoxia pathway, p53 pathway, PI3K, AKT, mTOR signaling and TNFα via NFκB signaling (Fig 4b, S10). Several pathways downregulated in SFTPC+ compared to SFTPC^−^/KRT17^low^ were also downregulated in SFTPC^−^/KRT17^low^ compared to KRT17^high^/KRT5^−^. These included EMT, inflammatory response, myogenesis and IL6/JAK/STAT3 pathway. Also, SFTPC^−^/KRT17^low^ compared to KRT17^high^/KRT5^−^ exhibit downregulation of Interferon α response and late Estrogen response (Fig 4c, S11).

**Fig. 4.**
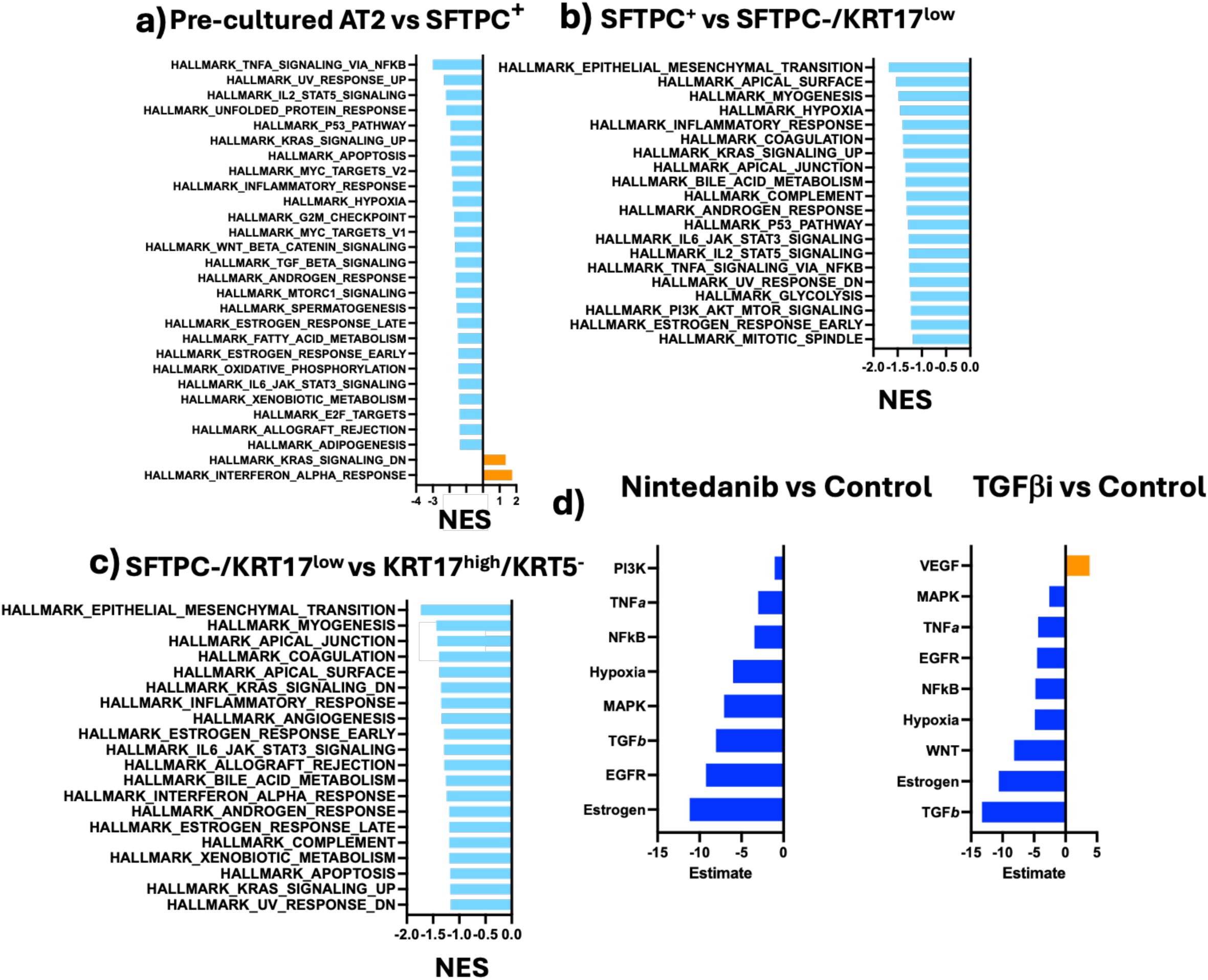
Pathways enriched in cultured epithelial cells. CD45^−^/EPCAM^+^/LysoTracker^high^ cells were co-cultured with MRC5 fibroblasts for 14 days and analyzed by single cell RNA sequencing. Enriched pathways were determined using Gene Set Enrichment Analysis (GSEA). Pathways significantly different (p ≤ 0.05) were arranged by the Normalized Enrichment Score (NES). Upregulated pathways (orange), downregulated (blue). **a)** Cultured SFTPC^+^ cells vs pre-cultured AT2 cells. **b)** SFTPC^+^ cells vs SFTPC^−^/KRT17^low^ cells. **c)** SFTPC^−^/KRT17^low^ cells vs. KRT17^high^/KRT5^−^. **d)** Estimated pathway activities from PROGENy showing significantly (FDR ≤ 0.1) down– and up-regulated signaling in epithelial cells after nintedanib or SB-431542 (TGFβi) treatment. Downregulated (blue), upregulated (orange).

PROGENy analyses was used to compare the effect of nintedanib and TGFβi on cultured epithelial cells, which leverage a large body of publicly available signaling perturbation experiments (46). There was an overlap between signaling pathway changes resulting from nintedanib and TGFβi treatments. Both downregulated TGFβ signaling, although the degree of downregulation was higher in TGFβi-treated cells. Both nintedanib and TGFβi also downregulated estrogen, hypoxia, NFκb, and TNFα pathways. While MAPK pathway was downregulated by both treatments, the degree of downregulation was higher in nintedanib-treated cells. PI3K signaling was significantly downregulated by nintedanib alone, while TGFβi downregulated the Wnt pathway and upregulated VEGF (Fig 4d).

### Nintedanib treatment induces significant gene expression changes in MRC5 fibroblasts

The effect nintedanib has on the differentiation of AT2 cells co-cultured with fibroblasts could occur directly or by modulating factors secreted by fibroblasts. To investigate the latter possibility, gene expression changes in MRC5 fibroblasts co-cultured with AT2 cells were quantified. Both nintedanib– and TGFβi-treated fibroblasts downregulated MTORC pathway, glycolysis and unfolded protein response. Both upregulated interferon α and γ response pathways (Fig 5a, S11, S12). There were also differences in the effect of nintedanib and TGFβi on fibroblast gene expression. Nintedanib, but not TGFβi, downregulated apoptosis, PI3K, protein secretion, and oxidative phosphorylation pathways (Fig 5a, S12). TGFβi, but not nintedanib, downregulated E2F target, mitotic spindle and G2/M checkpoint pathway, as well as upregulated IL6/JAK/STAT3 signaling (Fig. 5a, S13). TNFα signaling vs NFκB pathway was downregulated in nintedanib and upregulated in TGFβi-treated cells, suggesting that this pathway may play an opposite role at different stages of AT2 cell differentiation in culture. Nintedanib-treated fibroblasts also downregulated DNA repair and p53 signaling pathways (Fig. 5a). PROGENy analyses of nintedanib and TGFβi-treated fibroblast revealed less overlap than was found in epithelial cells: Both treatments upregulated JAK-STAT pathway. However, nintedanib alone downregulated MAPK, EGFR, PI3K and VEGF pathways, while TGFβi downregulated TGFβ pathway (Fig. 5b).

**Fig. 5.**
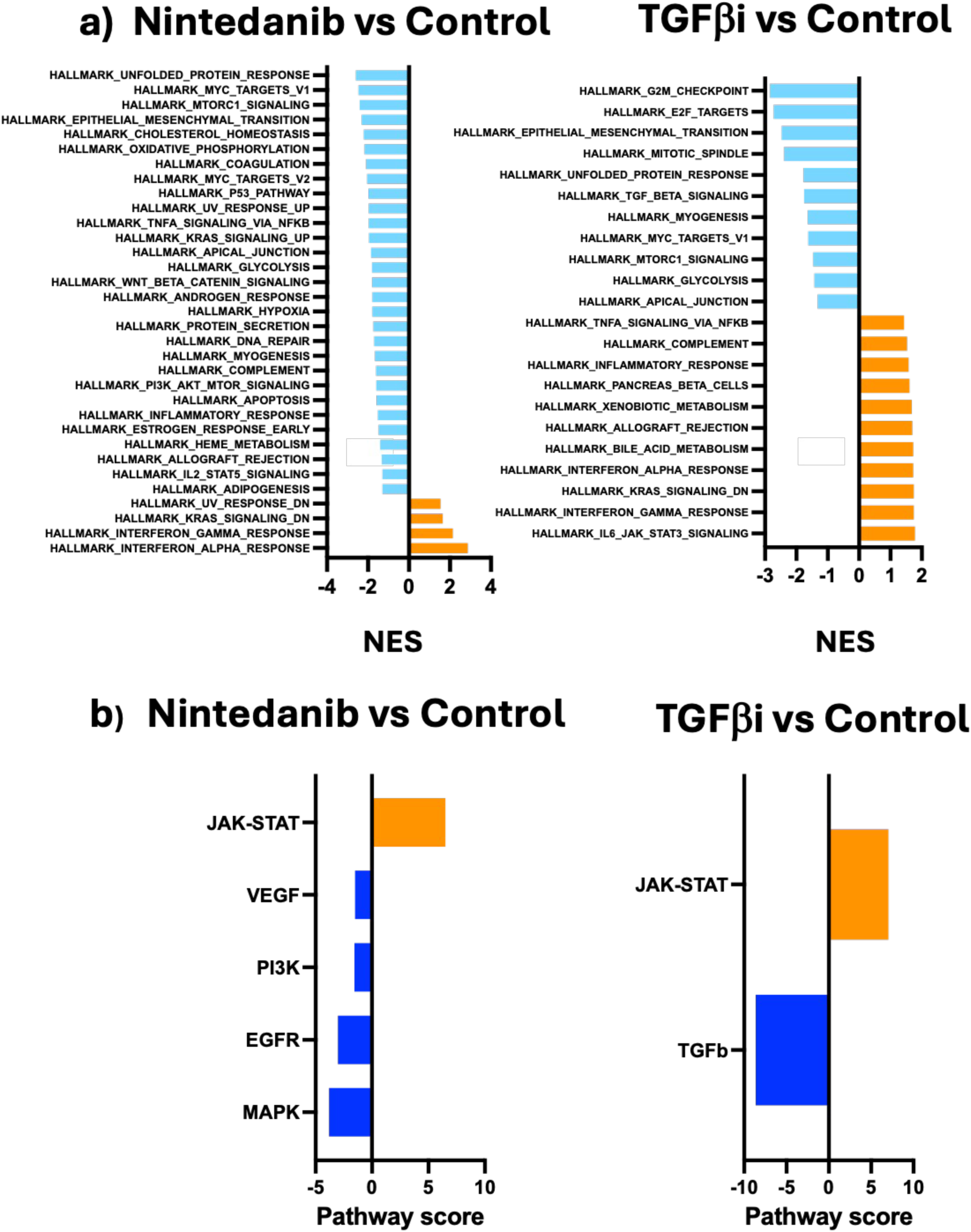
Pathway enrichment in Fibroblasts. CD45^−^/EPCAM^+^/LysoTracker^high^ cells were co-cultured with MRC5 fibroblasts for 14 days, treated as indicated and analyzed by scRNA-seq. **a)** Gene Set Enrichment Analysis (GSEA) of fibroblast transcriptomes co-cultures with epithelial cells: Pathways significantly (p ≤ 0.05) different were arranged by the Normalized Enrichment Score (NES). Upregulated pathways (orange), downregulated (blue). **b)** Estimated pathway activities from PROGENy showing significantly (FDR ≤ 0.1) down– and up-regulated signaling. Upregulated pathways (orange), downregulated (blue).

To further investigate whether the nintedanib effect on AT2 cells was mediated by fibroblasts, we interrogated the intercellular communication pathways between these cell types using the CellChat package (31), defining fibroblasts as senders and cultured epithelial cells as receivers (Fig. 6a). Compared to control, nintedanib treatment substantially altered the fibroblast secretome, enhancing signaling pathways involving pleiotrophin (*PTN*) and midkine (*MDK*), while suppressing multiple pro-fibrotic pathways, including those mediated by collagens, *PLAU*, *SEMA5A*, *POSTN*, *TNC*, *FN1*, and *AREG* (Fig. 6b). To dissect these effects further, we compared them to changes induced by TGFβi. While both treatments shared significant overlap, TGFβi induced a broader spectrum of upregulated ligand-receptor interactions (Fig. S14). A direct comparison of gene expression changes in fibroblast-secreted ligands revealed changes unique to nintedanib. *LIF*, *PLAU* and *SEMA5A* were uniquely downregulated in nintedanib-treated fibroblasts, while *PTN* was specifically upregulated (Fig. 6c-d). Furthermore, while both treatments enhanced *MDK* and suppressed *AREG*, these effects were more pronounced following nintedanib treatment (Fig 6c-d).

**Fig. 6.**
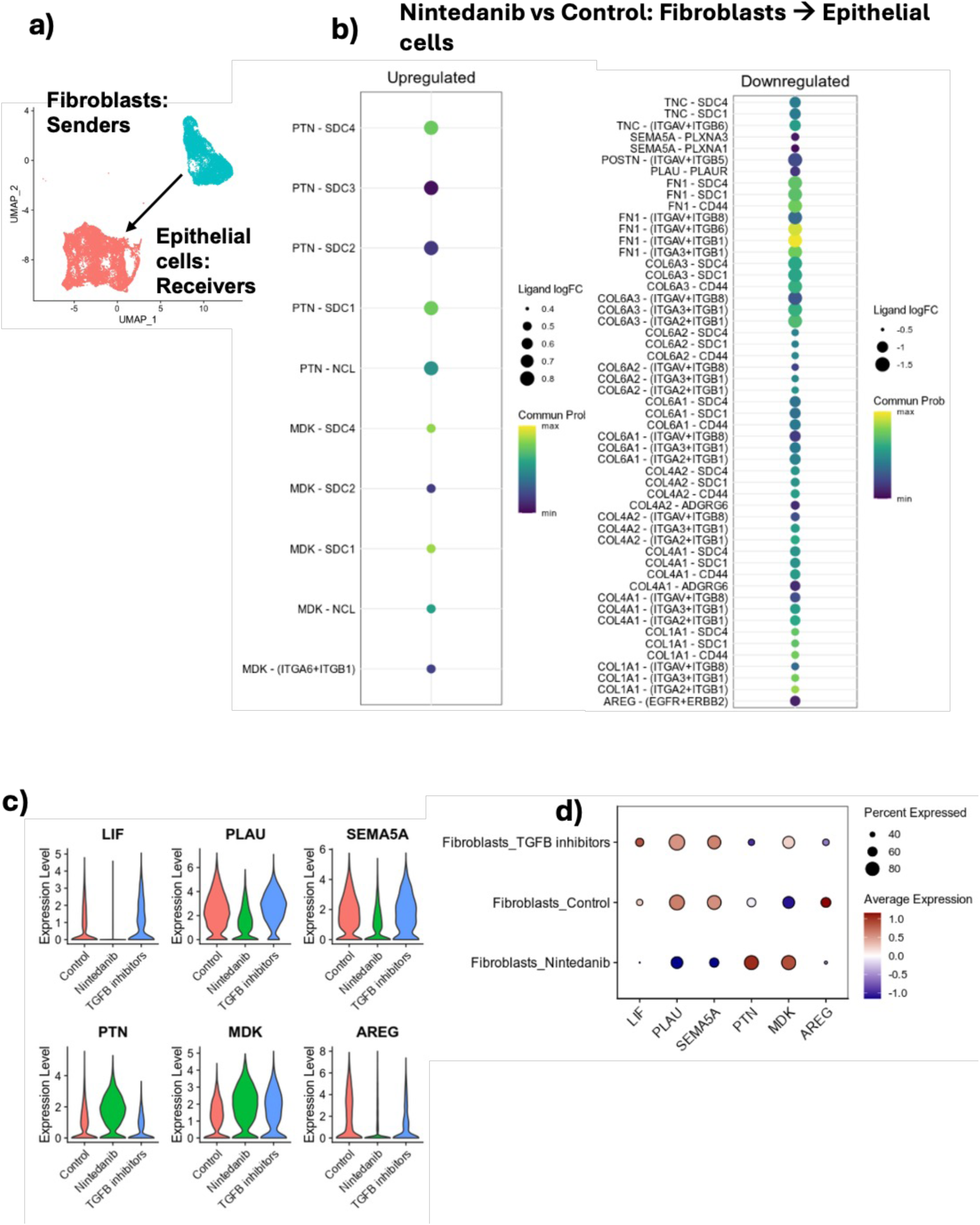
CellChat analyses of nintedanib treated AT2/MRC5 co-cultures. **a)** Schematic of CellChat analyses. **b)** Dot plots indicating the inferred communication probability of receptor-ligand pairs from upregulated (left panel) and downregulated (right panel) interaction pathways in nintedanib-treated cells compared to control. Violin plots **(c)** and dot plot **(d)** showing the expression levels of selected ligands in control, nintedanib– and TGFβi-treated fibroblasts.

To investigate if factors specifically affected by nintedanib in fibroblasts were responsible for the increased proliferation and AT2-like properties of cultured epithelial cells, IPF-AT2 cell organoids were treated with uPAi, LIFi and PTN. Unlike nintedanib treratment, none of these compounds improved colony formation (Fig S15), suggesting that nintedanib did not act via these factors individually.

## DISCUSSION

This study investigated whether the therapeutic efficacy of pirfenidone and nintedanib could in part be due to direct or indirect modulation of epithelial cell fate. The results showed that both pirfenidone and nintedanib significantly increase AT2 cell organoid formation. While control and pirfenidone-treated cells differentiate to abnormal basaloid cells, nintedanib-treated cells remain in an AT2-like state characterized by significant SFTPC expression. This is consistent with a previous report suggesting that nintedanib, but not pirfenidone stabilized SFTPC expression in both human and murine *ex vivo* lung cultures (47). Furthermore, effects of nintedanib are distinct from TGFβ inhibition, as AT2 cells treated with a TGFβ inhibitor lose SFTPC expression and are arrested in an intermediate state between the cells treated with nintedanib and untreated controls.

Here we report that after 14 days in culture, human AT2 cells differentiated to abnormal basaloid KRT17^high^/KRT5^−^ cells. However, a prior study had shown that AT2 cells co-cultured with adult human lung mesenchyme (AHLM) adopted a KRT5^+^ basal fate, while AT2 cells co-cultured with MRC5 fibroblasts retained AT2 cell characteristics (12). This discrepancy may reflect late passage-dependent changes in the secretome of the MRC5 fibroblasts used.

These data suggests a model where SFTPC^+^, AT2-like cells transition to SFTPC^−^/KRT17^low^ state, and then to KRT17^high^ state. Nintedanib increases the SFTPC^+^ state, blocking the progression to SFTPC^−^/KRT17^low^ while TGFβ inhibitor acts later, blocking the progression to the KRT17^high^ state, indicating that TGFβ promotes this state (Fig 7). From this it is possible that while there is some overlap between nintedanib and TGFβ signaling, nintedanib acts upstream of TGFβ during differentiation of AT2 cells. Also, AT2 cells co-cultured with AHLM differentiate to basal, KRT5^+^ cells (12), while our AT2 cells, co-cultured with late passage MRC5, stop at KRT17^high^/KRT5^−^ stage. This suggests that differentiation of abnormal basaloid to basal epithelial cells requires an additional factor that is present in AHML but absent in MRC5 fibroblasts.

**Fig. 7.**
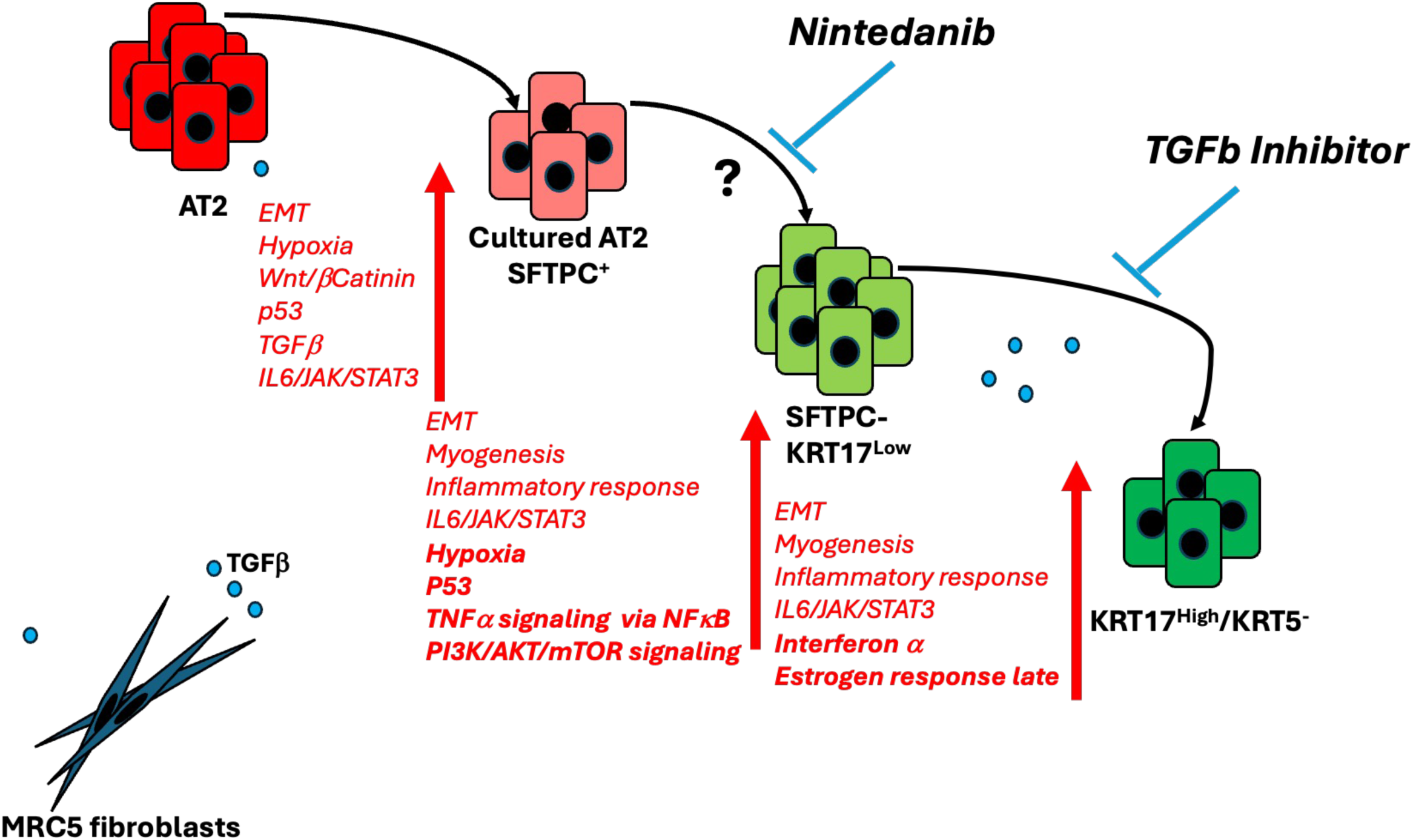
Working model of epithelial differentiation control by Nintedanib and TGFβ. Schematic summarizing the proposed trajectory from SFTPC^+^ AT2-like cells to KRT17^high^/KRT5^−^ basaloid cells, with nintedanib acting upstream of TGFβ.

Pathway analysis provides further insight. EMT signaling and IL6/JAK/STAT3 pathway are upregulated throughout progression from AT2 cells to KRT17^high^/KRT5^−^ abnormal basaloid stage (Fig 7) and have been implicated in IPF progression (48–50). This suggests that both nintedanib and TGFβi may act, in part, through suppressing these pathways. Indeed, EMT could be downregulated through a reduction of TGFβ signaling which also occurs in nintedanib-treated epithelial cells (51–54). Similarly, nintedanib has been shown to attenuate fibrosis in murine models through inhibiting JAK2 expression (55).

TNFα signaling via NFκB, PI3K/AKT/mTOR signaling and p53 pathway are upregulated earlier in AT2 cell transdifferentiation and are not changed during SFTPC^−^/KRT17^low^ progression to KRT17^high^/KRT5^−^ (Fig 7). Nintedanib has been reported to target both NFκB signaling and PI3K/AKT/mTOR pathway in rodent models (56, 57). Conversely, Interferon α and Estrogen response late pathway are involved in late stages of AT2 cell differentiation and may be regulated by TGFβ but not nintedanib (Fig 7).

The effect of pirfenidone and nintedanib on AT2 cells can be either direct or indirect through fibroblasts, and additional experiments are needed to differentiate these possibilities. Additionally, the organoid model used here represents only two cell types, AT2 cells and fibroblasts, and excludes endothelial and immune components that could also modulate epithelial cell fate and contribute to IPF progression (58, 59). More complex models, such as precision cut lung slices, may reveal how pirfenidone and nintedanib act on other cell types *in vivo*.

Although additional studies are required to understand the mechanism of action of nintedanib and pirfenidone, several clinically relevant conclusions can be drawn from these data: Both drugs act to increase AT2 organoid formation, consistent with improved AT2 cell fitness. This raises the possibility of using these medications to treat other lung diseases of AT2 cell failure, such as acute lung injury. Nintedanib treatment of three COVID-19 patients requiring intubation was followed by rapid improvement in lung function, suggesting that nintedanib may play a role in reversing COVID-19 damage to the lungs (60). Similarly, pirfenidone use was associated with a trend toward greater improved lung function following severe COVID infection (61). Additional testing of these findings in larger clinical trials is required to establish whether nintedanib or pirfenidone improve outcomes in these contexts. Finally, because pirfenidone and nintedanib act on AT2 cells via different mechanisms, patient responses may be heterogeneous. Clinical studies are required to define patient subgroups and potentially develop predictive biomarkers, allowing personalized therapy selection (62).

Here, we have demonstrated that nintedanib and pirfenidone enhance AT2 cell organoid formation and that nintedanib promotes an SFTPC^+^ AT2 cell phenotype, preventing their differentiation to abnormal basaloid cells via a mechanism that is distinct from TGFβ inhibition. Overall, these data suggest that an element of the “antifibrotic” activity of nintedanib may be due to preserving AT2 cell fate or functions, and that restoring AT2 cell function may be a therapeutic approach for managing IPF.

## REFERENCES

1. Mora AL, Rojas M, Pardo A, Selman M. Emerging therapies for idiopathic pulmonary fibrosis, a progressive age-related disease. Nat Rev Drug Discov. 2017;16(11):755–72.

2. Marshall DC, Salciccioli JD, Shea BS, Akuthota P. Trends in mortality from idiopathic pulmonary fibrosis in the European Union: an observational study of the WHO mortality database from 2001-2013. Eur Respir J. 2018;51(1).

3. Glass DS, Grossfeld D, Renna HA, Agarwala P, Spiegler P, Kasselman LJ, et al. Idiopathic pulmonary fibrosis: Molecular mechanisms and potential treatment approaches. Respir Investig. 2020;58(5):320–35.

4. Tomos I, Kanellopoulou P, Nastos D, Aidinis V. Pharmacological targeting of ECM homeostasis, fibroblast activation and invasion for the treatment of pulmonary fibrosis. Expert Opin Ther Targets. 2025;29(1-2):43–57.

5. Tsukui T, Wolters PJ, Sheppard D. Alveolar fibroblast lineage orchestrates lung inflammation and fibrosis. Nature. 2024;631(8021):627–34.

6. Parimon T, Yao C, Stripp BR, Noble PW, Chen P. Alveolar Epithelial Type II Cells as Drivers of Lung Fibrosis in Idiopathic Pulmonary Fibrosis. Int J Mol Sci. 2020;21(7).

7. Barkauskas CE, Cronce MJ, Rackley CR, Bowie EJ, Keene DR, Stripp BR, et al. Type 2 alveolar cells are stem cells in adult lung. J Clin Invest. 2013;123(7):3025–36.

8. Xu Y, Mizuno T, Sridharan A, Du Y, Guo M, Tang J, et al. Single-cell RNA sequencing identifies diverse roles of epithelial cells in idiopathic pulmonary fibrosis. JCI Insight. 2016;1(20):e90558.

9. Alder JK, Chen JJ, Lancaster L, Danoff S, Su SC, Cogan JD, et al. Short telomeres are a risk factor for idiopathic pulmonary fibrosis. Proc Natl Acad Sci U S A. 2008;105(35):13051–6.

10. Lee JS, La J, Aziz S, Dobrinskikh E, Brownell R, Jones KD, et al. Molecular markers of telomere dysfunction and senescence are common findings in the usual interstitial pneumonia pattern of lung fibrosis. Histopathology. 2021;79(1):67–76.

11. Chilosi M, Carloni A, Rossi A, Poletti V. Premature lung aging and cellular senescence in the pathogenesis of idiopathic pulmonary fibrosis and COPD/emphysema. Transl Res. 2013;162(3):156–73.

12. Kathiriya JJ, Wang C, Zhou M, Brumwell A, Cassandras M, Le Saux CJ, et al. Human alveolar type 2 epithelium transdifferentiates into metaplastic KRT5(+) basal cells. Nat Cell Biol. 2022;24(1):10–23.

13. Noble PW, Albera C, Bradford WZ, Costabel U, Glassberg MK, Kardatzke D, et al. Pirfenidone in patients with idiopathic pulmonary fibrosis (CAPACITY): two randomised trials. Lancet. 2011;377(9779):1760–9.

14. King TE, Jr., Bradford WZ, Castro-Bernardini S, Fagan EA, Glaspole I, Glassberg MK, et al. A phase 3 trial of pirfenidone in patients with idiopathic pulmonary fibrosis. N Engl J Med. 2014;370(22):2083–92.

15. Noble PW, Albera C, Bradford WZ, Costabel U, du Bois RM, Fagan EA, et al. Pirfenidone for idiopathic pulmonary fibrosis: analysis of pooled data from three multinational phase 3 trials. Eur Respir J. 2016;47(1):243–53.

16. Nathan SD, Albera C, Bradford WZ, Costabel U, Glaspole I, Glassberg MK, et al. Effect of pirfenidone on mortality: pooled analyses and meta-analyses of clinical trials in idiopathic pulmonary fibrosis. Lancet Respir Med. 2017;5(1):33–41.

17. Richeldi L, du Bois RM, Raghu G, Azuma A, Brown KK, Costabel U, et al. Efficacy and safety of nintedanib in idiopathic pulmonary fibrosis. N Engl J Med. 2014;370(22):2071–82.

18. Flaherty KR, Wells AU, Cottin V, Devaraj A, Inoue Y, Richeldi L, et al. Nintedanib in progressive interstitial lung diseases: data from the whole INBUILD trial. Eur Respir J. 2022;59(3).

19. Varone F, Sgalla G, Iovene B, Bruni T, Richeldi L. Nintedanib for the treatment of idiopathic pulmonary fibrosis. Expert Opin Pharmacother. 2018;19(2):167–75.

20. Torre A, Martinez-Sanchez FD, Narvaez-Chavez SM, Herrera-Islas MA, Aguilar-Salinas CA, Cordova-Gallardo J. Pirfenidone use in fibrotic diseases: What do we know so far? Immun Inflamm Dis. 2024;12(7):e1335.

21. Ying H, Fang M, Hang QQ, Chen Y, Qian X, Chen M. Pirfenidone modulates macrophage polarization and ameliorates radiation-induced lung fibrosis by inhibiting the TGF-beta1/Smad3 pathway. J Cell Mol Med. 2021;25(18):8662–75.

22. Ma HY, Vander Heiden JA, Uttarwar S, Xi Y, N’Diaye EN, LaCanna R, et al. Inhibition of MRTF activation as a clinically achievable anti-fibrotic mechanism for pirfenidone. Eur Respir J. 2023;61(4).

23. C L, A C, L V, L B, d’Alessandro M, P C, et al. Common molecular pathways targeted by nintedanib in cancer and IPF: A bioinformatic study. Pulm Pharmacol Ther. 2020;64:101941.

24. Van der Velden JL, Bertoncello I, McQualter JL. LysoTracker is a marker of differentiated alveolar type II cells. Respir Res. 2013;14(1):123.

25. Yun J, Hansen S, Morris O, Madden DT, Libeu CP, Kumar AJ, et al. Senescent cells perturb intestinal stem cell differentiation through Ptk7 induced noncanonical Wnt and YAP signaling. Nature Communications. 2023;14(1):156.

26. Liberzon A, Birger C, Thorvaldsdóttir H, Ghandi M, Mesirov JP, Tamayo P. The Molecular Signatures Database (MSigDB) hallmark gene set collection. Cell Syst. 2015;1(6):417–25.

27. Korotkevich G, Sukhov V, Budin N, Shpak B, Artyomov MN, Sergushichev A. Fast gene set enrichment analysis. bioRxiv. 2021:060012.

28. Wu D, Smyth GK. Camera: a competitive gene set test accounting for inter-gene correlation. Nucleic Acids Research. 2012;40(17):e133-e.

29. Holland CH, Tanevski J, Perales-Patón J, Gleixner J, Kumar MP, Mereu E, et al. Robustness and applicability of transcription factor and pathway analysis tools on single-cell RNA-seq data. Genome Biology. 2020;21(1):36.

30. Badia-i-Mompel P, Vélez Santiago J, Braunger J, Geiss C, Dimitrov D, Müller-Dott S, et al. decoupleR: ensemble of computational methods to infer biological activities from omics data. Bioinformatics Advances. 2022;2(1).

31. Jin S, Guerrero-Juarez CF, Zhang L, Chang I, Ramos R, Kuan CH, et al. Inference and analysis of cell-cell communication using CellChat. Nat Commun. 2021;12(1):1088.

32. Street K, Risso D, Fletcher RB, Das D, Ngai J, Yosef N, et al. Slingshot: cell lineage and pseudotime inference for single-cell transcriptomics. BMC Genomics. 2018;19(1):477.

33. Zheng SC, Stein-O’Brien G, Augustin JJ, Slosberg J, Carosso GA, Winer B, et al. Universal prediction of cell-cycle position using transfer learning. Genome Biol. 2022;23(1):41.

34. Jacobs JP, Jones CM, Baille JP. Characteristics of a human diploid cell designated MRC-5. Nature. 1970;227(5254):168–70.

35. Fernandez IE, Eickelberg O. The impact of TGF-beta on lung fibrosis: from targeting to biomarkers. Proc Am Thorac Soc. 2012;9(3):111–6.

36. Tatler AL, Jenkins G. TGF-beta activation and lung fibrosis. Proc Am Thorac Soc. 2012;9(3):130–6.

37. Rangarajan S, Kurundkar A, Kurundkar D, Bernard K, Sanders YY, Ding Q, et al. Novel Mechanisms for the Antifibrotic Action of Nintedanib. Am J Respir Cell Mol Biol. 2016;54(1):51–9.

38. Jin J, Togo S, Kadoya K, Tulafu M, Namba Y, Iwai M, et al. Pirfenidone attenuates lung fibrotic fibroblast responses to transforming growth factor-beta1. Respir Res. 2019;20(1):119.

39. Inman GJ, Nicolas FJ, Callahan JF, Harling JD, Gaster LM, Reith AD, et al. SB-431542 is a potent and specific inhibitor of transforming growth factor-beta superfamily type I activin receptor-like kinase (ALK) receptors ALK4, ALK5, and ALK7. Mol Pharmacol. 2002;62(1):65–74.

40. Cai XT, Jia M, Heigl T, Shamir ER, Wong AK, Hall BM, et al. IL-4-induced SOX9 confers lineage plasticity to aged adult lung stem cells. Cell Reports. 2024;43(8).

41. Jennings P, Bertocchi C, Frick M, Haller T, Pfaller W, Dietl P. Ca2+ induced surfactant secretion in alveolar type II cultures isolated from the H-2Kb-tsA58 transgenic mouse. Cell Physiol Biochem. 2005;15(1-4):159–66.

42. Zacharias WJ, Frank DB, Zepp JA, Morley MP, Alkhaleel FA, Kong J, et al. Regeneration of the lung alveolus by an evolutionarily conserved epithelial progenitor. Nature. 2018;555(7695):251–5.

43. Adams TS, Schupp JC, Poli S, Ayaub EA, Neumark N, Ahangari F, et al. Single-cell RNA-seq reveals ectopic and aberrant lung-resident cell populations in idiopathic pulmonary fibrosis. Sci Adv. 2020;6(28):eaba1983.

44. Habermann AC, Gutierrez AJ, Bui LT, Yahn SL, Winters NI, Calvi CL, et al. Single-cell RNA sequencing reveals profibrotic roles of distinct epithelial and mesenchymal lineages in pulmonary fibrosis. Sci Adv. 2020;6(28):eaba1972.

45. Gold MP, Reyes M, Diamant N, Kuo T, Hajiramezanali E, Newburger JW, et al. Foundation Model Attributions Reveal Shared Inflammatory Program Across Diseases. bioRxiv. 2025:2025.06.14.659567.

46. Schubert M, Klinger B, Klunemann M, Sieber A, Uhlitz F, Sauer S, et al. Perturbation-response genes reveal signaling footprints in cancer gene expression. Nat Commun. 2018;9(1):20.

47. Lehmann M, Buhl L, Alsafadi HN, Klee S, Hermann S, Mutze K, et al. Differential effects of Nintedanib and Pirfenidone on lung alveolar epithelial cell function in ex vivo murine and human lung tissue cultures of pulmonary fibrosis. Respir Res. 2018;19(1):175.

48. Hill C, Jones MG, Davies DE, Wang Y. Epithelial-mesenchymal transition contributes to pulmonary fibrosis via aberrant epithelial/fibroblastic cross-talk. J Lung Health Dis. 2019;3(2):31–5.

49. Kim KK, Kugler MC, Wolters PJ, Robillard L, Galvez MG, Brumwell AN, et al. Alveolar epithelial cell mesenchymal transition develops in vivo during pulmonary fibrosis and is regulated by the extracellular matrix. Proc Natl Acad Sci U S A. 2006;103(35):13180–5.

50. Montero P, Milara J, Roger I, Cortijo J. Role of JAK/STAT in Interstitial Lung Diseases; Molecular and Cellular Mechanisms. Int J Mol Sci. 2021;22(12).

51. Willis BC, Liebler JM, Luby-Phelps K, Nicholson AG, Crandall ED, du Bois RM, et al. Induction of epithelial-mesenchymal transition in alveolar epithelial cells by transforming growth factor-beta1: potential role in idiopathic pulmonary fibrosis. Am J Pathol. 2005;166(5):1321–32.

52. Willis BC, Borok Z. TGF-beta-induced EMT: mechanisms and implications for fibrotic lung disease. Am J Physiol Lung Cell Mol Physiol. 2007;293(3):L525–34.

53. Ihara H, Mitsuishi Y, Kato M, Takahashi F, Tajima K, Hayashi T, et al. Nintedanib inhibits epithelial-mesenchymal transition in A549 alveolar epithelial cells through regulation of the TGF-beta/Smad pathway. Respir Investig. 2020;58(4):275–84.

54. Hughes D, Prestle J, Zippel N, McFetridge S, Szczepan M, Neubauer H, et al. Nintedanib Induces Mesenchymal-to-Epithelial Transition and Reduces Subretinal Fibrosis Through Metabolic Reprogramming. Int J Mol Sci. 2025;26(15).

55. Yang Y, Wang X, Zhang J. Pirfenidone and nintedanib attenuate pulmonary fibrosis in mice by inhibiting the expression of JAK2. J Thorac Dis. 2024;16(2):1128–40.

56. Hao R, Li Y, Li J, Guo Z, Yang Z, Lu W. Nintedanib alleviates hyperoxia-induced lung injury via targeting NF-kappaB signalling pathway in rat model of bronchopulmonary dysplasia. Folia Histochem Cytobiol. 2025;63(2):79–87.

57. Pan L, Cheng Y, Yang W, Wu X, Zhu H, Hu M, et al. Nintedanib Ameliorates Bleomycin-Induced Pulmonary Fibrosis, Inflammation, Apoptosis, and Oxidative Stress by Modulating PI3K/Akt/mTOR Pathway in Mice. Inflammation. 2023;46(4):1531–42.

58. Mutsaers SE, Miles T, Prele CM, Hoyne GF. Emerging role of immune cells as drivers of pulmonary fibrosis. Pharmacol Ther. 2023;252:108562.

59. Bian F, Lan YW, Zhao S, Deng Z, Shukla S, Acharya A, et al. Lung endothelial cells regulate pulmonary fibrosis through FOXF1/R-Ras signaling. Nat Commun. 2023;14(1):2560.

60. Bussolari C, Palumbo D, Fominsky E, Nardelli P, De Lorenzo R, Vitali G, et al. Case Report: Nintedaninb May Accelerate Lung Recovery in Critical Coronavirus Disease 2019. Front Med (Lausanne). 2021;8:766486.

61. Bermudo-Peloche G, Del Rio B, Vicens-Zygmunt V, Bordas-Martinez J, Hernandez M, Valenzuela C, et al. Pirfenidone in post-COVID-19 pulmonary fibrosis (FIBRO-COVID): a phase 2 randomised clinical trial. Eur Respir J. 2025;65(4).

62. Wick KD, Ware LB, Matthay MA. Acute respiratory distress syndrome. Bmj. 2024;387:e076612.

