## Supplemental Figures for "Nintedanib and Pirfenidone Affect Growth and Differentiation of Human Alveolar Type 2 Cells"

### Slide 1
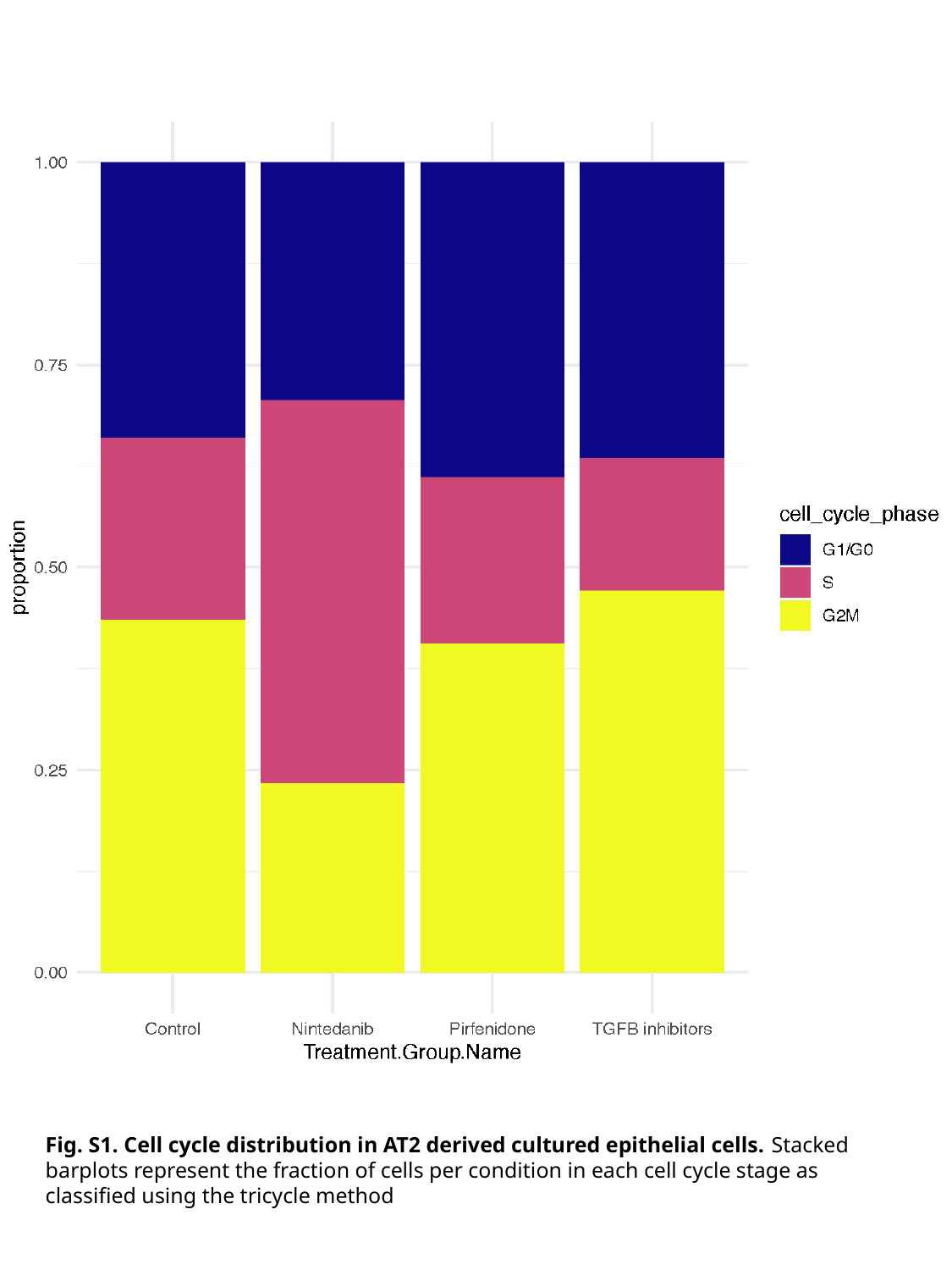

Fig. S1. Cell cycle distribution in AT2 derived cultured epithelial cells. Stacked barplots represent the fraction of cells per condition in each cell cycle stage as classified using the tricycle method

### Slide 2
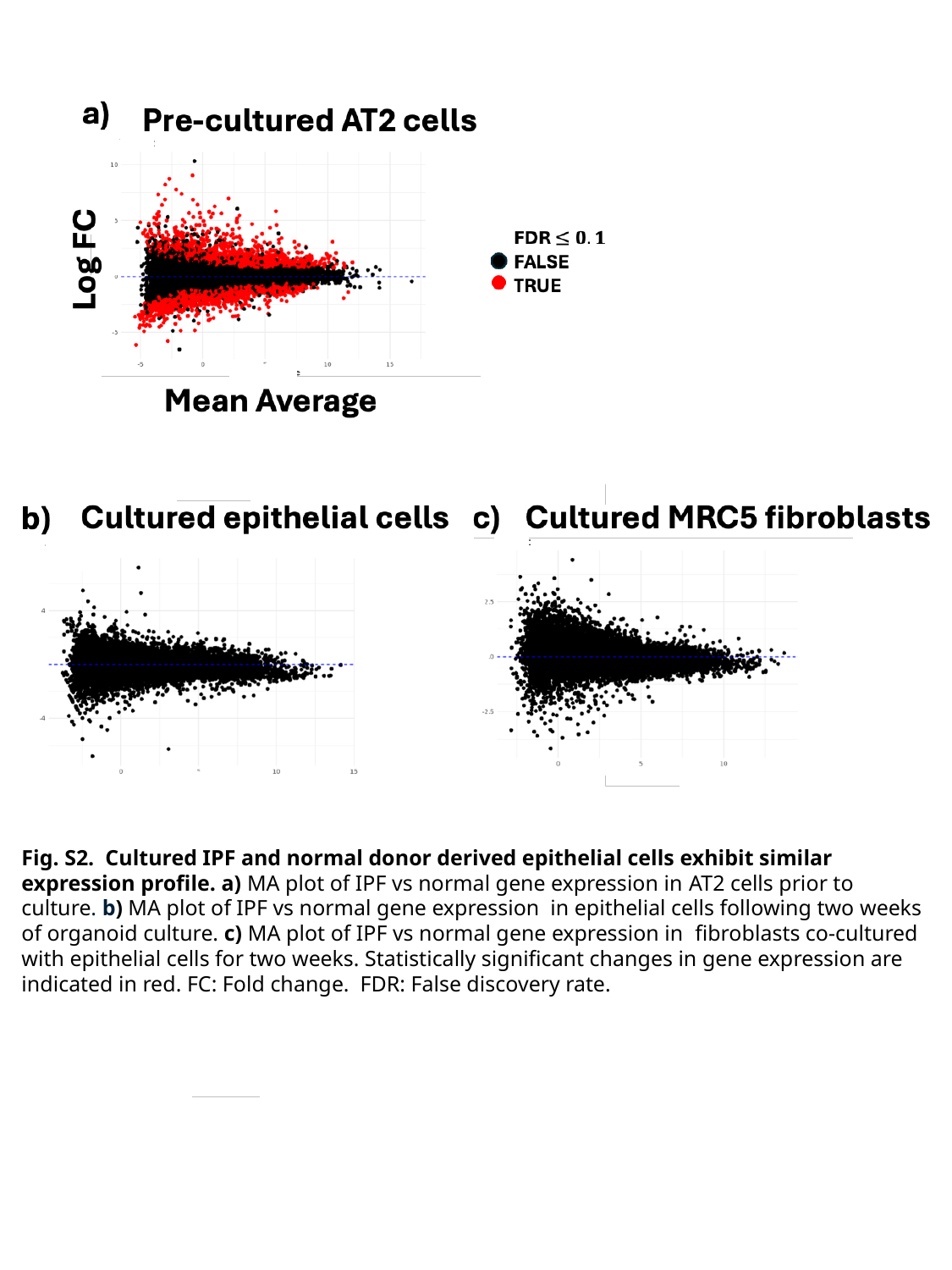

Fig. S2. Cultured IPF and normal donor derived epithelial cells exhibit similar expression profile. a) MA plot of IPF vs normal gene expression in AT2 cells prior to culture. b) MA plot of IPF vs normal gene expression in epithelial cells following two weeks of organoid culture. c) MA plot of IPF vs normal gene expression in fibroblasts co-cultured with epithelial cells for two weeks. Statistically significant changes in gene expression are indicated in red. FC: Fold change. FDR: False discovery rate.

### Slide 3
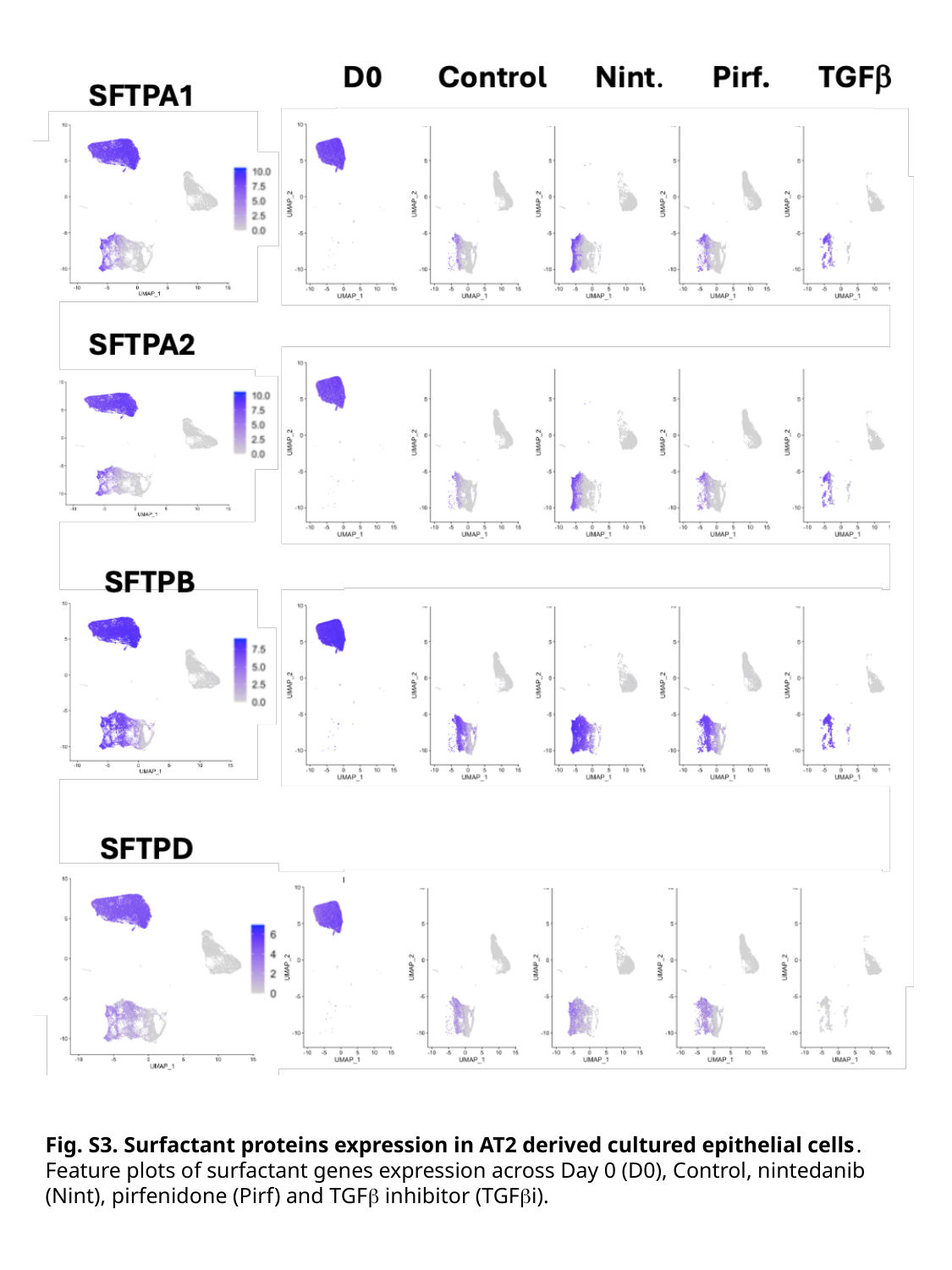

Fig. S3. Surfactant proteins expression in AT2 derived cultured epithelial cells. Feature plots of surfactant genes expression across Day 0 (D0), Control, nintedanib (Nint), pirfenidone (Pirf) and TGFβ inhibitor (TGFβi).

### Slide 4
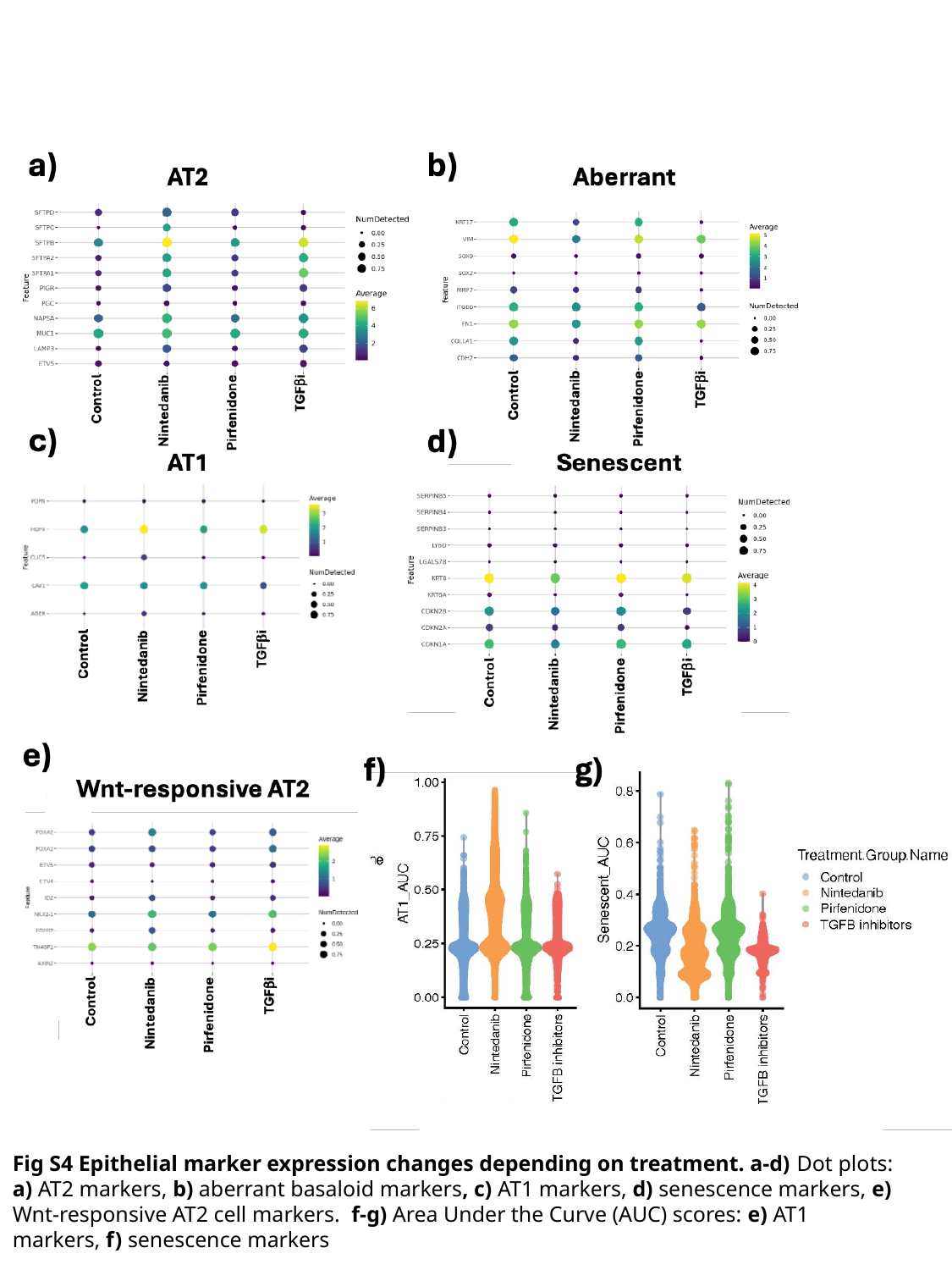

Fig S4 Epithelial marker expression changes depending on treatment. a-d) Dot plots: a) AT2 markers, b) aberrant basaloid markers, c) AT1 markers, d) senescence markers, e) Wnt-responsive AT2 cell markers. f-g) Area Under the Curve (AUC) scores: e) AT1 markers, f) senescence markers

### Slide 5
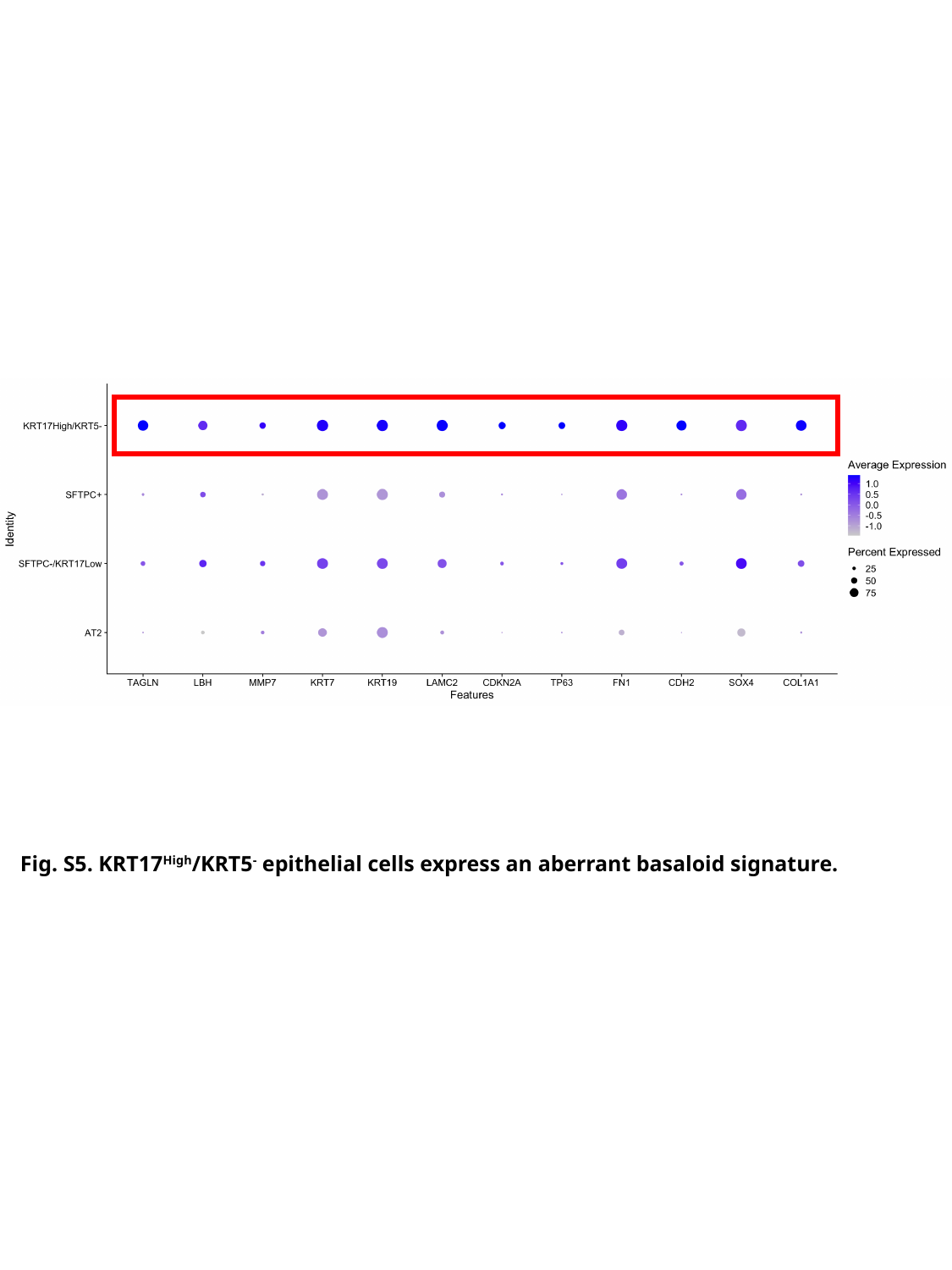

Fig. S5. KRT17^High^/KRT5^-^ epithelial cells express an aberrant basaloid signature.

### Slide 6
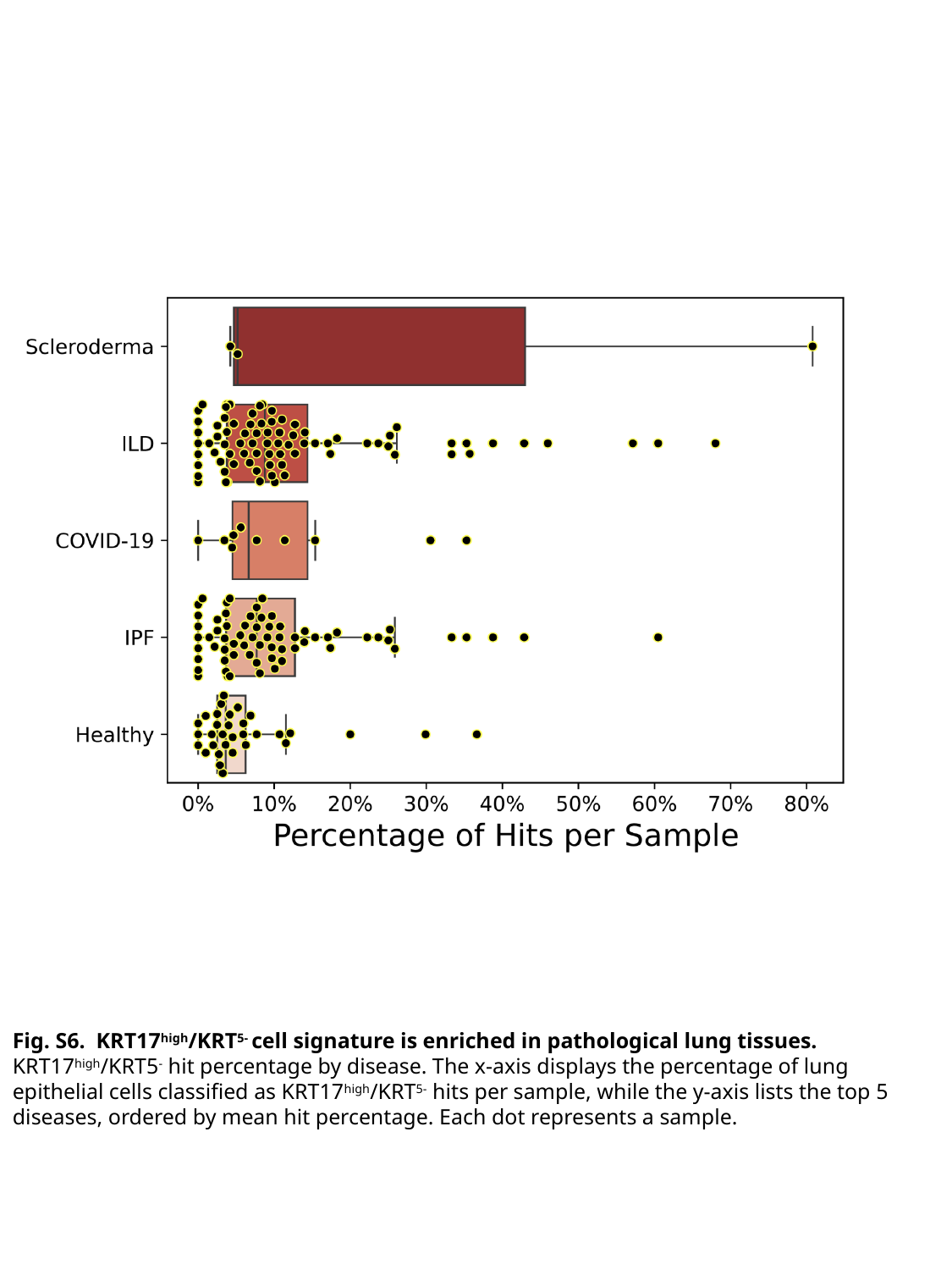

Fig. S6. KRT17^high^/KRT^5-^ cell signature is enriched in pathological lung tissues. KRT17^high^/KRT5^-^ hit percentage by disease. The x-axis displays the percentage of lung epithelial cells classified as KRT17^high^/KRT^5-^ hits per sample, while the y-axis lists the top 5 diseases, ordered by mean hit percentage. Each dot represents a sample.

### Slide 7
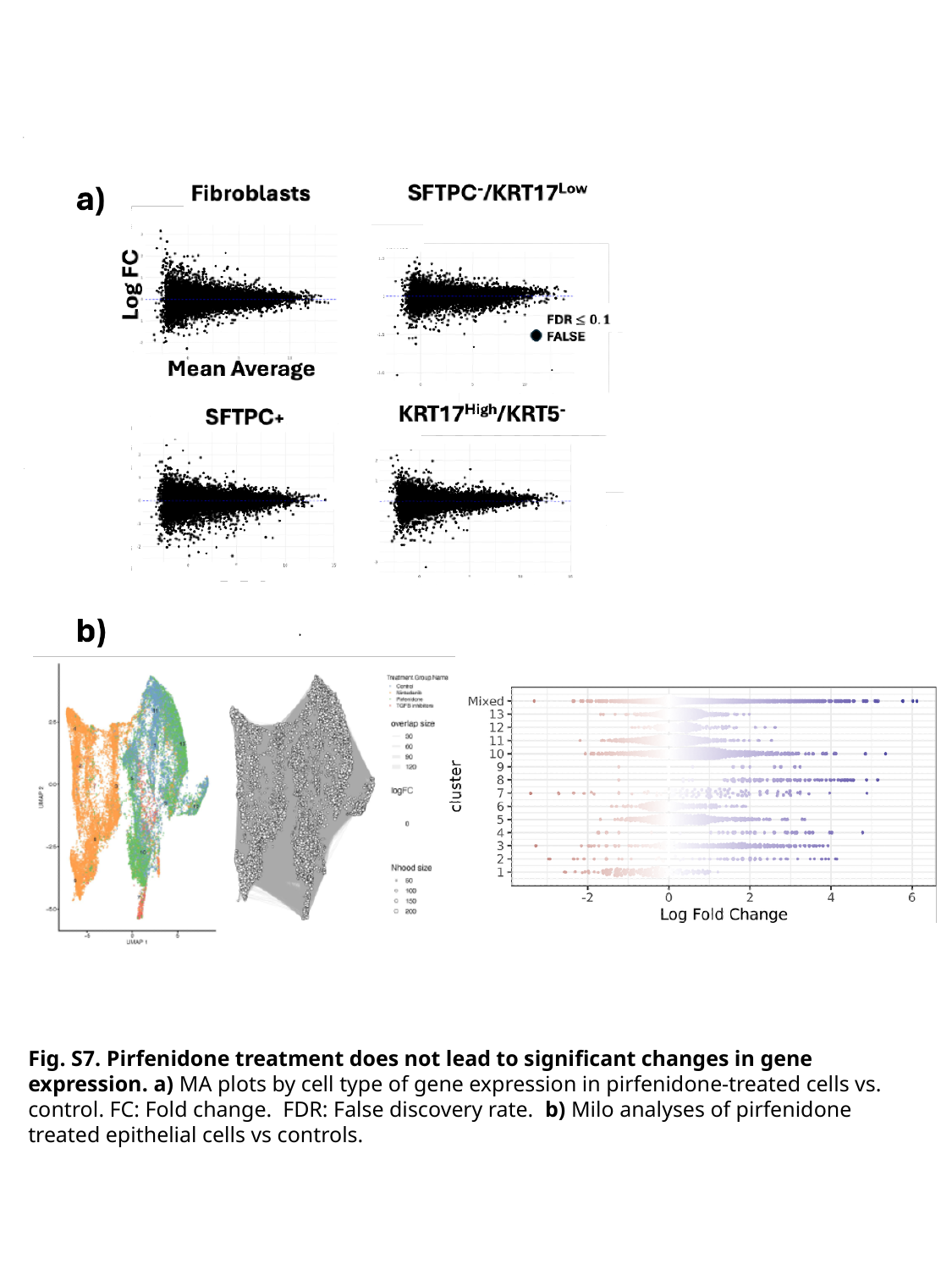

Fig. S7. Pirfenidone treatment does not lead to significant changes in gene expression. a) MA plots by cell type of gene expression in pirfenidone-treated cells vs. control. FC: Fold change. FDR: False discovery rate. b) Milo analyses of pirfenidone treated epithelial cells vs controls.

### Slide 8
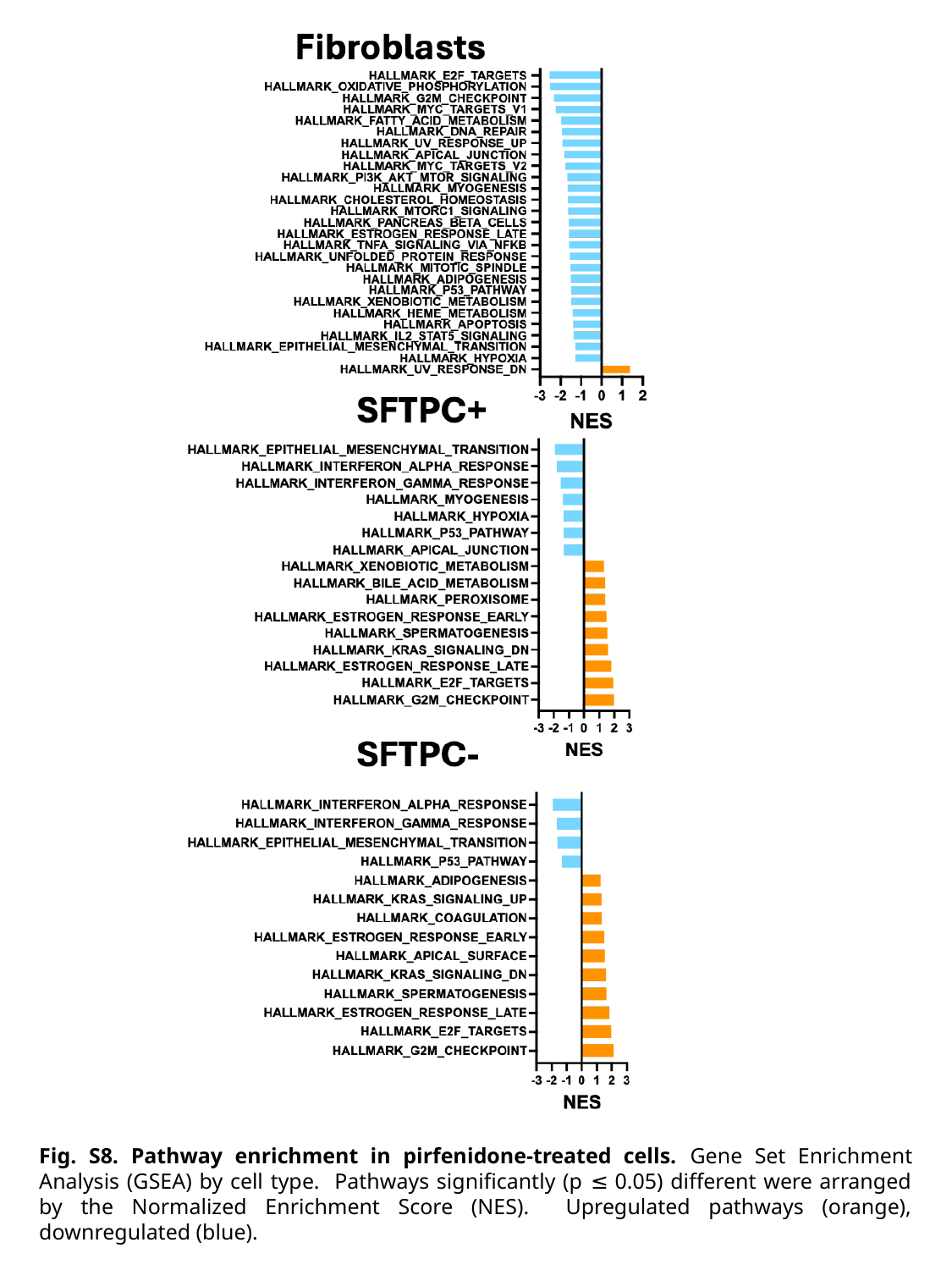

Fig. S8. Pathway enrichment in pirfenidone-treated cells. Gene Set Enrichment Analysis (GSEA) by cell type. Pathways significantly (p ≤ 0.05) different were arranged by the Normalized Enrichment Score (NES). Upregulated pathways (orange), downregulated (blue).

### Slide 9
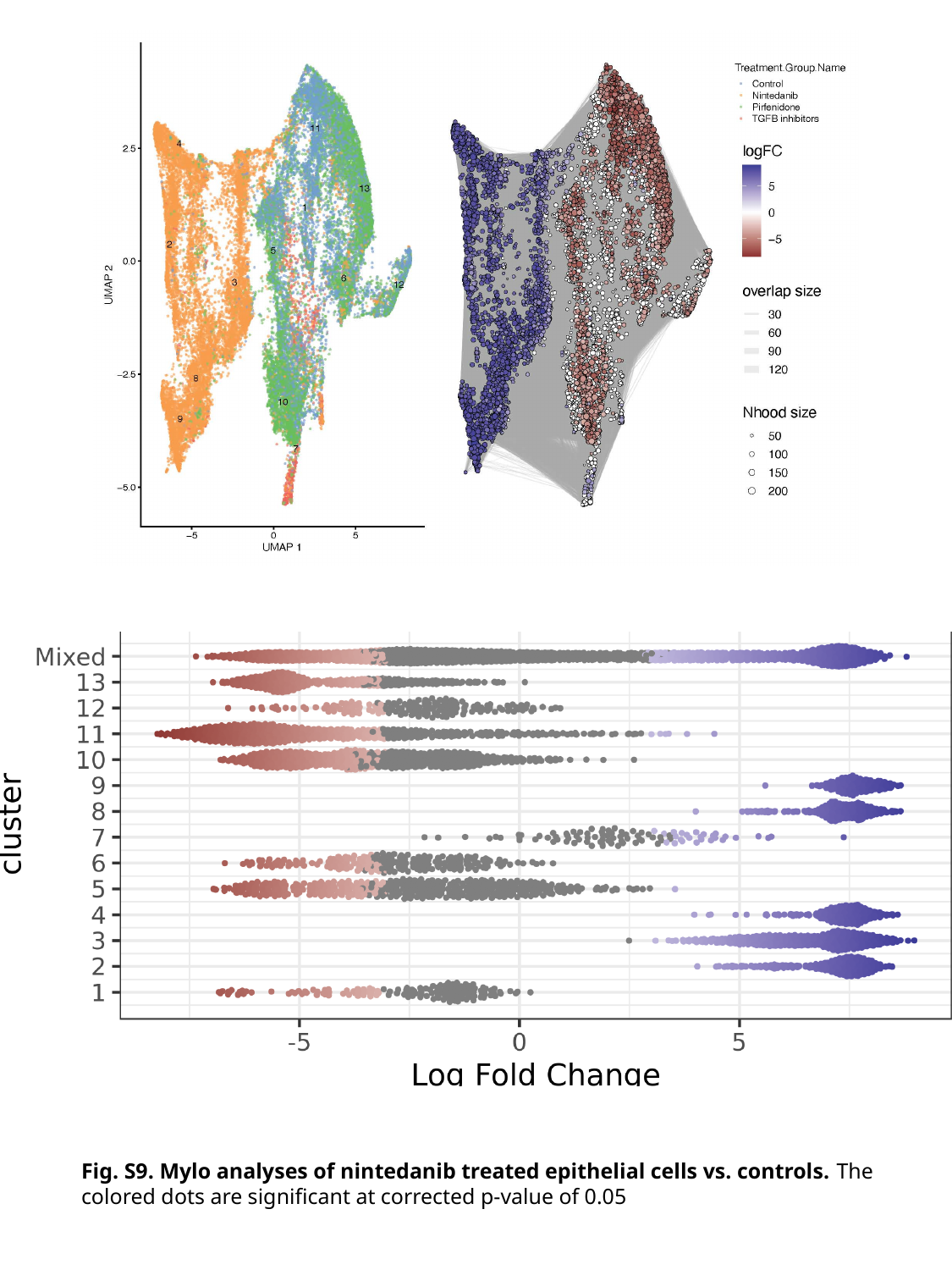

Fig. S9. Mylo analyses of nintedanib treated epithelial cells vs. controls. The colored dots are significant at corrected p-value of 0.05

### Slide 10
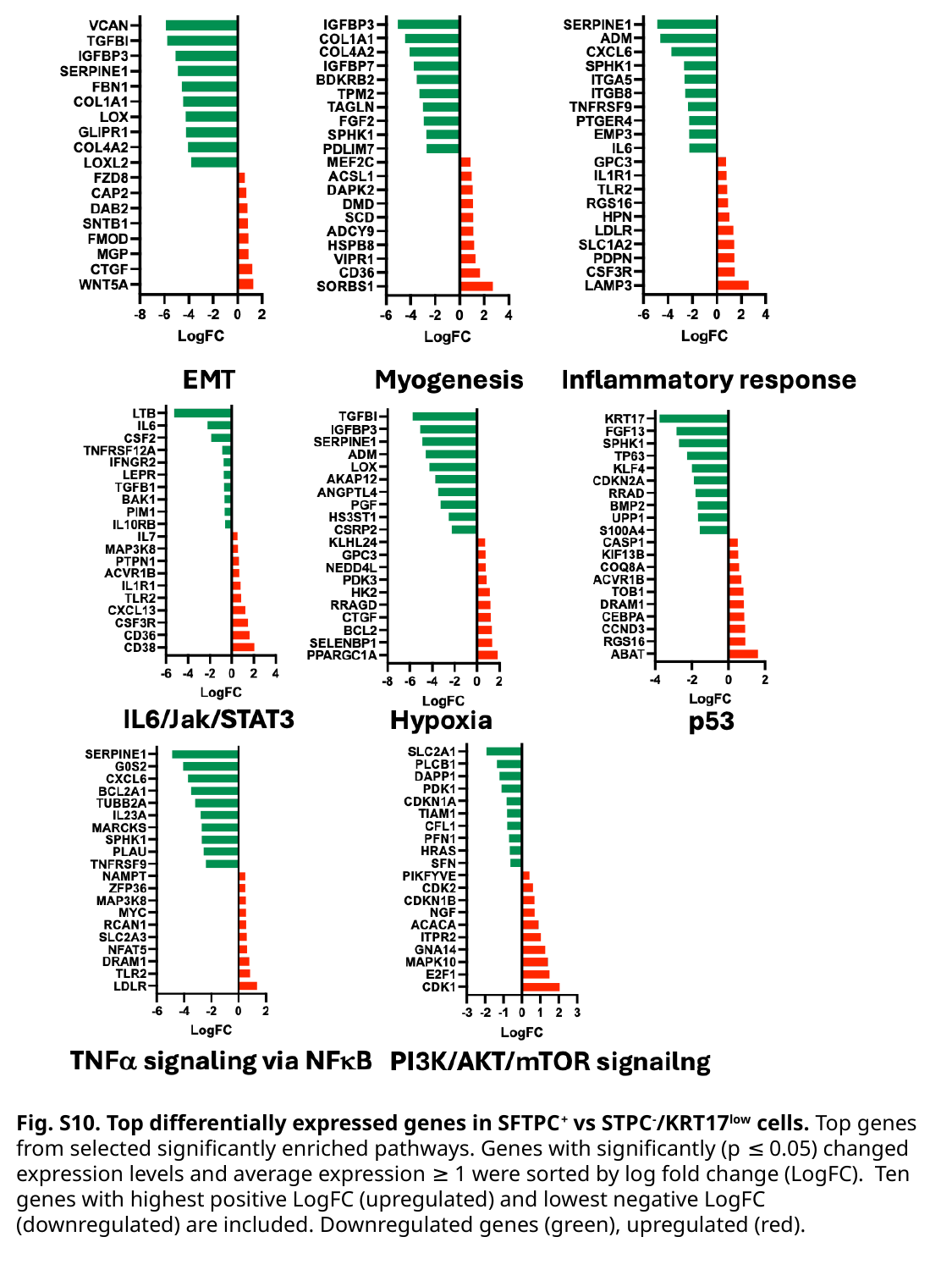

Fig. S10. Top differentially expressed genes in SFTPC^+^ vs STPC^-^/KRT17^low^ cells. Top genes from selected significantly enriched pathways. Genes with significantly (p ≤ 0.05) changed expression levels and average expression ≥ 1 were sorted by log fold change (LogFC). Ten genes with highest positive LogFC (upregulated) and lowest negative LogFC (downregulated) are included. Downregulated genes (green), upregulated (red).

### Slide 11
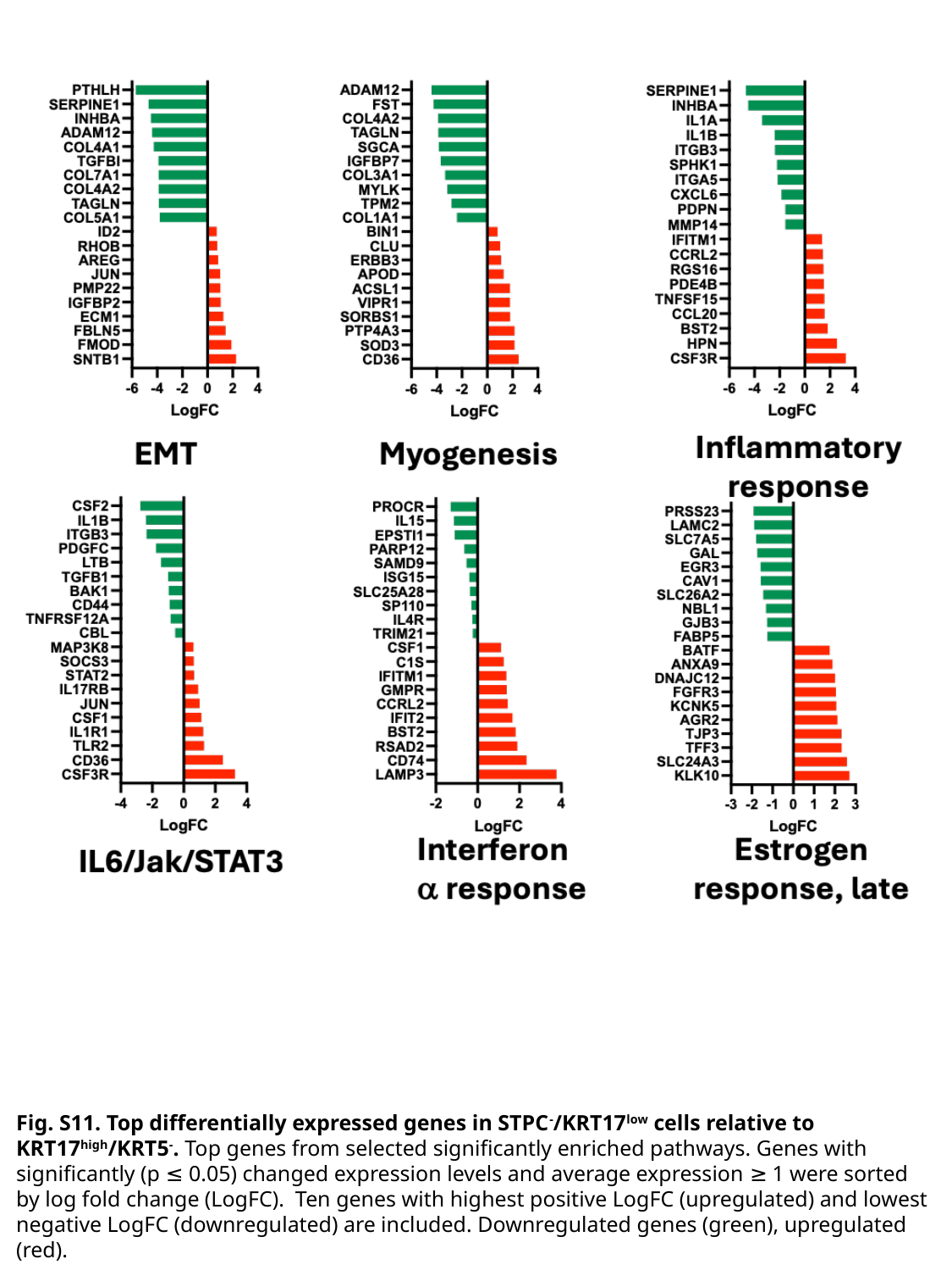

Fig. S11. Top differentially expressed genes in STPC^-^/KRT17^low^ cells relative to KRT17^high^/KRT5^-^. Top genes from selected significantly enriched pathways. Genes with significantly (p ≤ 0.05) changed expression levels and average expression ≥ 1 were sorted by log fold change (LogFC). Ten genes with highest positive LogFC (upregulated) and lowest negative LogFC (downregulated) are included. Downregulated genes (green), upregulated (red).

### Slide 12
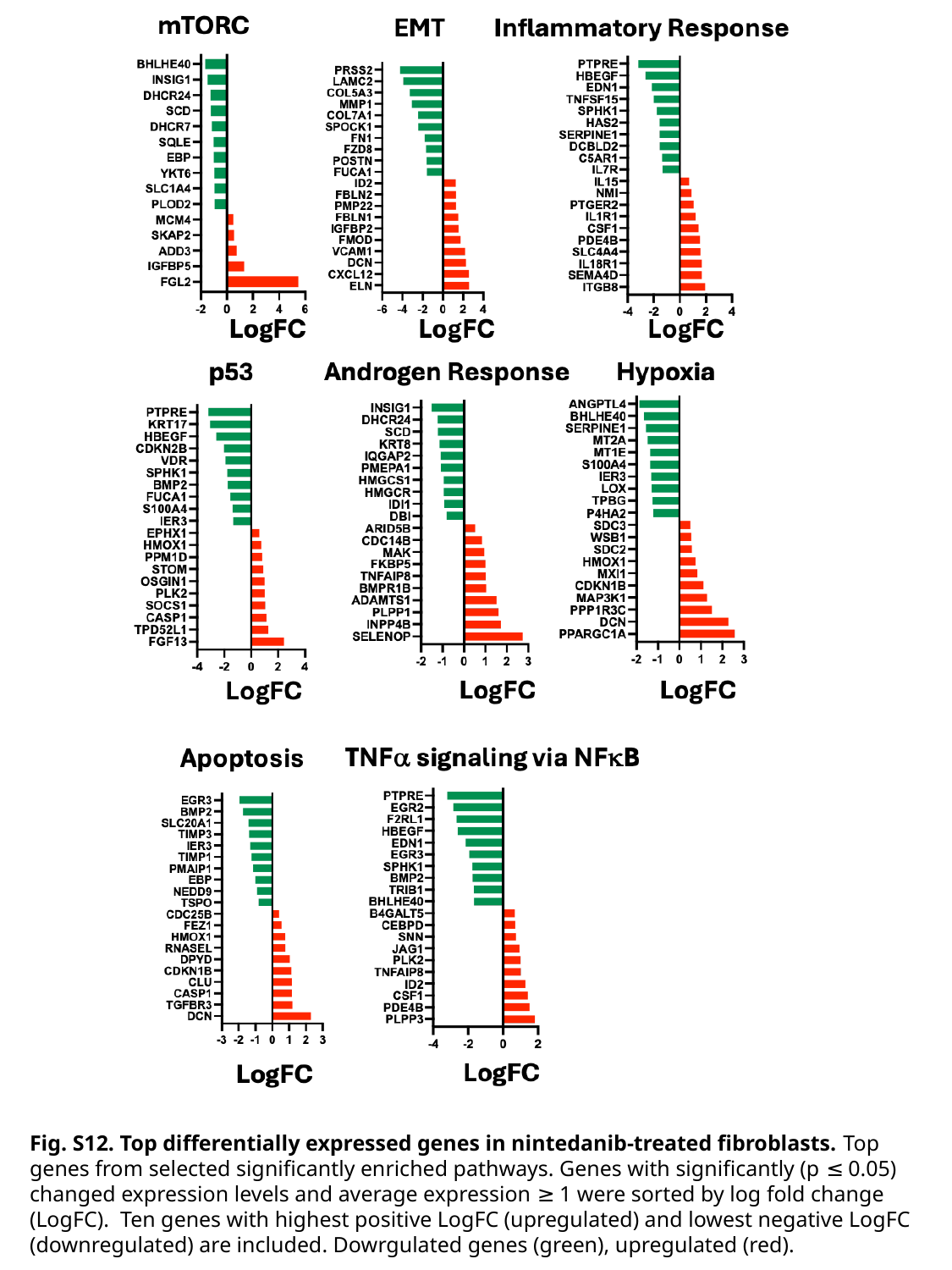

Fig. S12. Top differentially expressed genes in nintedanib-treated fibroblasts. Top genes from selected significantly enriched pathways. Genes with significantly (p ≤ 0.05) changed expression levels and average expression ≥ 1 were sorted by log fold change (LogFC). Ten genes with highest positive LogFC (upregulated) and lowest negative LogFC (downregulated) are included. Dowrgulated genes (green), upregulated (red).

### Slide 13
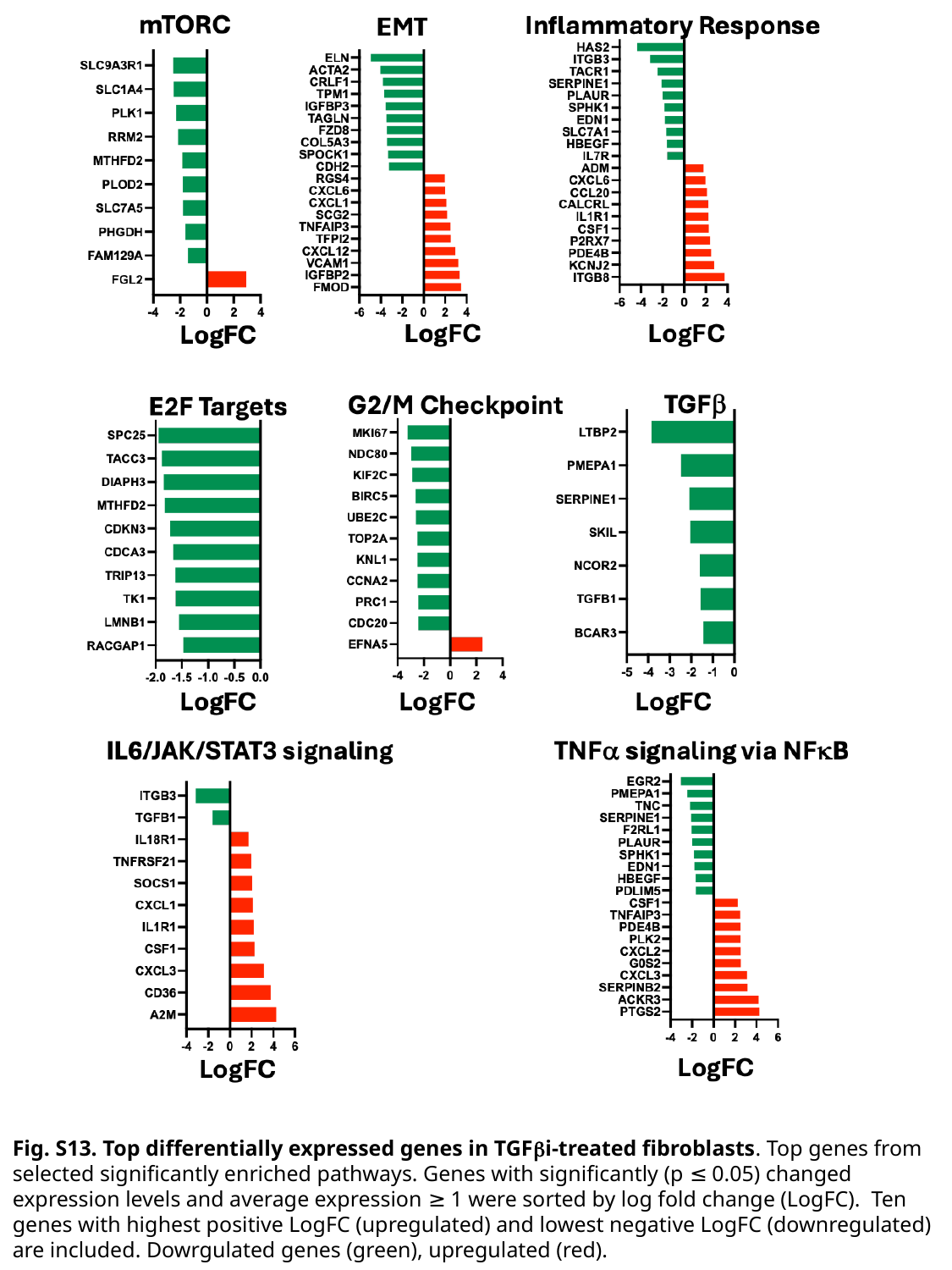

Fig. S13. Top differentially expressed genes in TGFβi-treated fibroblasts. Top genes from selected significantly enriched pathways. Genes with significantly (p ≤ 0.05) changed expression levels and average expression ≥ 1 were sorted by log fold change (LogFC). Ten genes with highest positive LogFC (upregulated) and lowest negative LogFC (downregulated) are included. Dowrgulated genes (green), upregulated (red).

### Slide 14
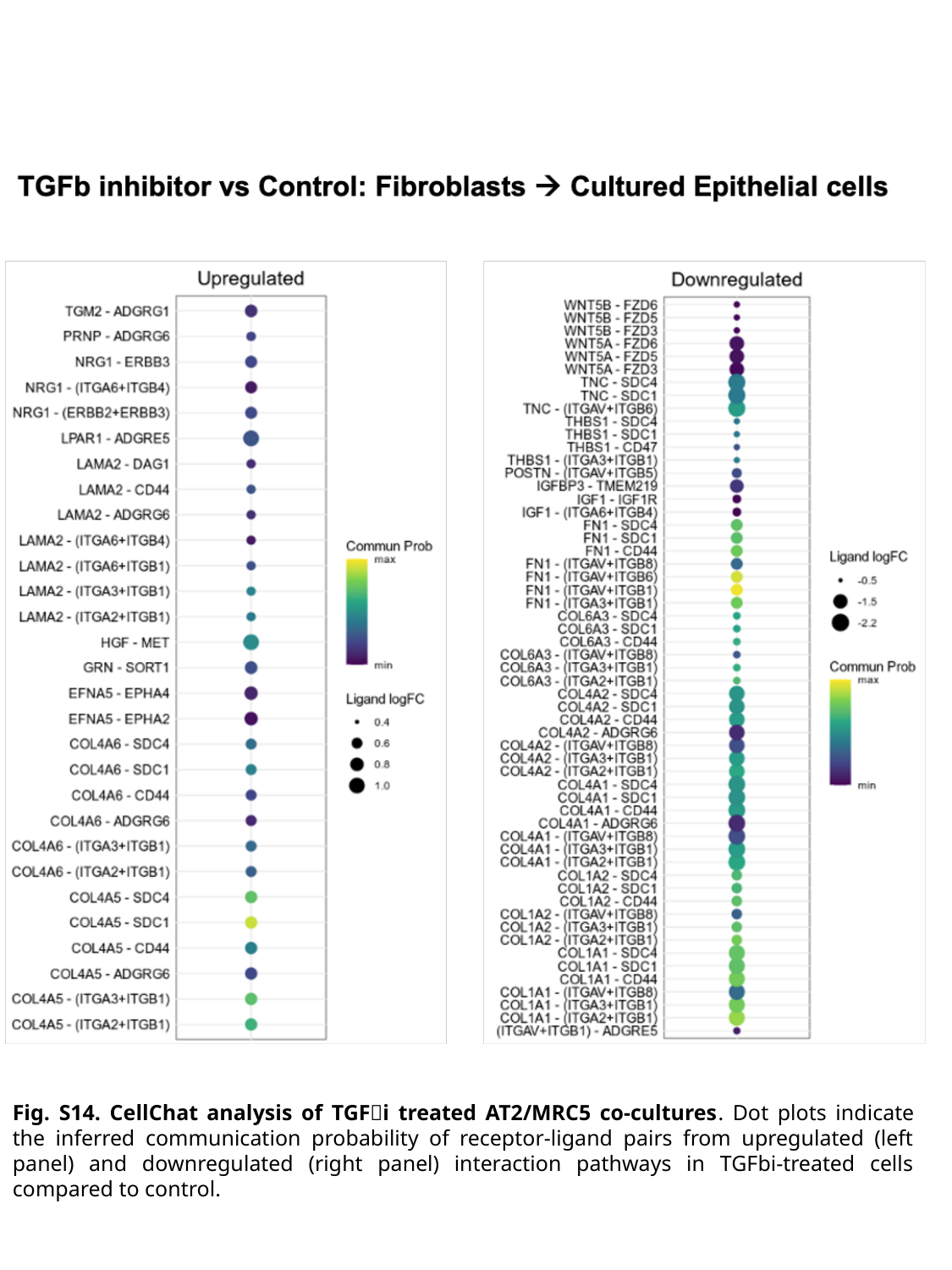

Fig. S14. CellChat analysis of TGFi treated AT2/MRC5 co-cultures. Dot plots indicate the inferred communication probability of receptor-ligand pairs from upregulated (left panel) and downregulated (right panel) interaction pathways in TGFbi-treated cells compared to control.

### Slide 15
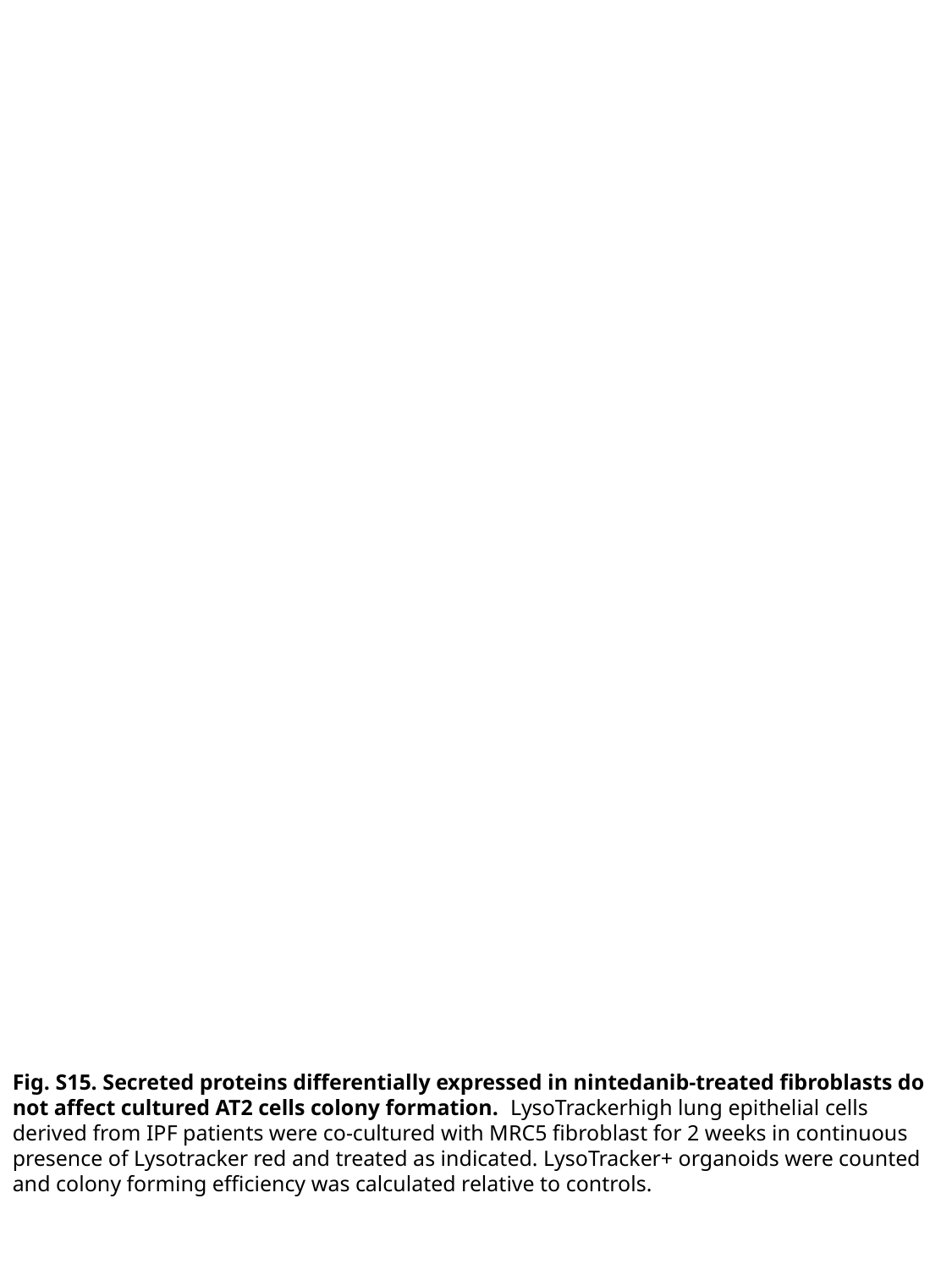

Fig. S15. Secreted proteins differentially expressed in nintedanib-treated fibroblasts do not affect cultured AT2 cells colony formation. LysoTrackerhigh lung epithelial cells derived from IPF patients were co-cultured with MRC5 fibroblast for 2 weeks in continuous presence of Lysotracker red and treated as indicated. LysoTracker+ organoids were counted and colony forming efficiency was calculated relative to controls.
