## Supplemental Methods and Table 1 for "Nintedanib and Pirfenidone Affect Growth and Differentiation of Human Alveolar Type 2 Cells"

**SUPPLEMENTAL INFORMATION**

**Supplemental Methods**

**Reagents:** Reagents were purchased from following vendors: pirfenidone (*TCI America, Portland, OR*, # P1871), nintedanib (*MedChemExpress, Monmouth Junction, NJ*, #HY-50940), hPTN (*R&D systems, Minneapolis, MN*, #252-PL), TGFβi (SB-341542, *Cayman Chemical Company, Ann Arbor, MI*, #13031), uPAi (BC11 hydrobromide, *TORCIS, Bristol, United Kingdom*, #4372), LIFi (EC359, *MedChemExpress, Monmouth Junction, NJ*, #HY-120142).

**Isolation of human alveolar type 2 epithelial cells:** Human lungs were perfused with heparin in PBS and digested with Dispase (*Gibco*, *Waltham, MA)* (5U/mL) in DMEM for 60 min at 37°C. DNase, type IV (*Sigma*, *St. Louis, MO*) was added to the lung digest to the final concentration of 0.25 mg/ml. The tissue was minced further and serially filtered through 70 um and 40 um filters. Lung homogenate was suspended slowly on 30 % percoll with 70 % percoll (*Cytiva, Uppsala, Sweden*) made in PBS placed at the bottom of the tube. Centrifugation of lung homogenate and percoll gradient was carried out at 616g for 20 min. The top layer of supernatant, including the interphase, consisting of epithelial cells and immune cells was removed and centrifuged at 373g for 8 min. The supernatant was discarded, and the cell pellet was washed 2 X with PBS and resuspended in DMEM supplemented with Penicillin, Streptomycin and Fungizone. IgG plates were prepared by coating petri dishes with 0.5 mg/ml human IgG (*Equitech-Bio, Kerrville, TX*) in 50mM TRIS pH 9.5 for 3 hours and washed twice with PBS. Five ml/plate of lung cell homogenate was added to each petri dish and incubated for 30 min. The unattached cells were collected and centrifuged at 373 g for 5 min, washed in PBS twice followed by suspension in TEC+ media (1).

**Flow cytometry:** Human lung cell suspension was stained with LysoTracker red (*Invitrogen, Carlsbad, CA*) (1:20,000) in a tissue culture incubator at 37°C for 45 min, washed in PBS, resuspended in FACS buffer (PBS, 0.5% Bovine Serum Albumin) and stained with conjugated primary antibodies to EpCAM (9C4) (*BioLegend,* *San Diego, CA,* #324212) and CD45 (HI40) (*BioLegend,* *San Diego, CA,* #304022) for 45 min at +4 °C. The cells were washed with PBS and resuspended in FACS buffer (PBS, 0.5% BSA). Dead cell stain Sytox™ Blue (*Invitrogen, Carlsbad, CA*) was added prior to analyses. Cells were analyzed and sorted on BD Aria cell sorter (*Franklin Lakes, NJ*). Data was analyzed on FlowJo V10.

**TEC basic medium composition:** 1M Hepes 7.5ml, 200mM Glutamine 10ml, 7.5% NaHCO3 2 mL, Fungizone (1000X, 250 ug/ml) 500ul, Pencillin-streptomycin (100X, 10,000U or ug/ml) 5ml. Add DMEM/F-12 50:50 mix to a volume of 500 mL. The media prepared was sterile filtered, stored at 4^0^C and used within 4 weeks.

**TEC plus media composition:** To 234.45 mL of TEC basic media, following reagents were added to make TEC plus. Insulin (10ug/ml) 250ul, Transferrin (5ug/ml) 250ul, Cholera Toxin (0.1ug/ml final) 25ul, Epidermal growth factor (25ng/ml) 25ul, Bovine Pituitary extract (15mg/500ml) 2.5ml, FBS (5% final) 12.5ml. The media prepared was sterile filtered, stored at 4^0^C and used within 2 weeks.

**Immunofluorescence**: 4 uM sections of paraformaldehyde fixed, OCT (*Fisher HealthCare, Houston, TX*) embedded organoids suspended in matrigel were equilibrated to room temperature and subjected to antigen retrieval: Slides were microwaved in Na Citrate buffer (*Sigma, St Louis, MO*) for 6 min. and allowed to cool for 30 min. The slides were then washed in PBS (3X, 5 min) and permeabilized with 0.5% Triton X in PBS for 10 min. The slides were washed in PBS (3X, 5 min) and blocked in 3% BSA, 10% Donkey serum 0.1% Triton in PBS for 1 hour at room temperature. The slides were then incubated overnight with primary antibodies: anti-E-cadherin (goat, 1:100, AF648, *R&D Systems, Minneapolis, MN*); anti-Prosurfactant protein C (rabbit, 1:500, AB3786, *EMD Millipore,* *Burlington, MA*); anti-cytokeratin 17 (rabbit, 1:500, 17516-1-1AP, *Proteintech, Rosemont, IL*) in blocking solution at +4°C. Following washing with PBS, 0.1% Tween 20 (PBS-T) (3X, 5 min), the slides were incubated with secondary antibodies Alexa Fleur 594 donkey anti-rabbit and Alexa Fleur 488 donkey anti-goat (1:500, *BioLegend, San Diego, CA*) in blocking buffer for 1 hr. at room temperature. The slides were then washed in PBS-T (3x, 5 min) and mounted with VECTASHIELD Antifade Mounting Medium (*Vector Laboratories, Newark, CA*). The slides were imaged with Axio Imager M2 fluorescent microscope (*Zeiss, Oberkochen, Germany*) at 40X magnification.

**Harvesting cells from Matrigel:** Inserts with organoids embedded in Matrigel were transferred into fresh 24-well pate. 200μl Dispase (5U/ml) in PBS was added to each insert and the plate was incubated for 1 hr. at 37°C. Organoids were dislodged by pipetting up and down and removed into 15ml tube. The inserts were washed with additional 200μl Dispase. Organoids were spun down at 373g for 5 min, washed in PBS twice, resuspended in 1ml 0.05% Trypsin and incubated at 37°C for 2 min. Trypsin was inactivated with 10ml TEC+ media, the cells were centrifuged at 373g, resuspended in 2ml TEC+ and counted. For single cell RNA sequencing (scRNA-seq) cell concentration was adjusted to 1,200 cells/μl.

**ScRNA-seq alignment and processing:** After processing with the 10x chromium, the resulting libraries were sequenced on an Illumnia NovaSeq machine with the obtained images being processed to Fastq files using the bcl2fastq program following standard process. Fastq files were then aligned to hg38 (IGIS4) using CellRanger 9.0 with the include introns option set to True. To remove ambient RNA contamination frequently seen in scRNA-seq data that include AT2 cells, we further processed the CellRanger alignments using CellBender v 0.3.0 using default settings. Initial QC filtering as well as batch integration and clustering were performed in R using the scran.chan package. In brief, cells were initially removed from any sample if they were outside three median absolute deviations in terms of total number of reads, number of features detected, or fraction of reads mapping to mitochondrial genes. Upon visual inspection of the results, we did an additional filtering round, removing cells with fewer than 3162 total reads, 1000 genes, or more than 20% mitochondrial reads.

Following QC filtering, read counts were log transformed and size factors were calculated using the function logNormCounts.chan with batch set to Sample. To perform downstream PCA and batch integration, we selected the top 2000 most highly variable genes using the modelGeneVar.chan function with batch set to Sample followed by Principal Component Analysis on these genes using sample-weighted PCA in the function runPCA.chan. Batch correction of these PCs was then performed using fastMNN (2) as implemented in mnnCorrect.chan with neighborhood parameter k set to 15 and otherwise default parameters.

To identify clusters in our data, and to generate tSNE and UMAP embeddings, we processed our fastMNN corrected PCs using the runAllDownstream function from scran.chan using default parameters including k = 10, number of neighbors = 10, and method set to “multilevel” (Louvain) for clustering. Markers per cluster were calculated using the scoreMarkers function with markers ranked by AUC. Clusters were then given approximate cell type names based on the expression of these marker as well as visual inspection of key marker genes.

**Milo analyses.** To assess differences in cell type proportions between drug treatment and control cells in our cell culture system, we made use of the Milo package (3) as applied to our cultured epithelial cell types. Briefly, cultured epithelial cells were subset from our full single-cell dataset and re-integrated/re-clustered using the same scran.chan workflow as applied to the whole dataset. We then constructed a k-nearest neighbors graph using the buildGraph function with k = 40 using the first 25 dimensions of the MNN-corrected principal components. We assigned cells to overlapping neighborhoods using this knn graph using the same k and d values as for knn graph construction with a random sampling proportion of 0.3. Changes in cell type proportion were assessed with a linear model in limma (4) followed by spatial FDR correction as implemented in the Milo package.

**SIGnature analyses**. The SIGnature package (5) was used to query the KRT17high/KRT5- cells across human diseases in the Scimilarity dataset (6). The gene signature was composed of the top 30 differentially expressed genes relative to other epithelial cells identified using the *FindMarkers* function from Seurat (only.pos = T, min.pct = 0.5, logfc.threshold =1) (7). The SIGnature search was limited to lung epithelial cells for this analysis.

**Supplemental Table**

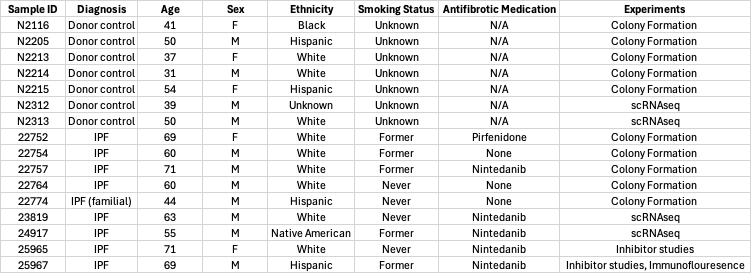

**Table S1. Patient demographics.**

**Supplemental References**

1.	Rowe RK, Brody SL, Pekosz A. Differentiated cultures of primary hamster tracheal airway epithelial cells. In Vitro Cell Dev Biol Anim. 2004;40(10):303-11.

2.	Haghverdi L, Lun ATL, Morgan MD, Marioni JC. Batch effects in single-cell RNA-sequencing data are corrected by matching mutual nearest neighbors. Nat Biotechnol. 2018;36(5):421-7.

3.	Dann E, Henderson NC, Teichmann SA, Morgan MD, Marioni JC. Differential abundance testing on single-cell data using k-nearest neighbor graphs. Nature Biotechnology. 2022;40(2):245-53.

4.	Ritchie ME, Phipson B, Wu D, Hu Y, Law CW, Shi W, et al. limma powers differential expression analyses for RNA-sequencing and microarray studies. Nucleic Acids Res. 2015;43(7):e47.

5.	Gold MP, Reyes M, Diamant N, Kuo T, Hajiramezanali E, Newburger JW, et al. Foundation Model Attributions Reveal Shared Inflammatory Program Across Diseases. bioRxiv. 2025:2025.06.14.659567.

6.	Heimberg G, Kuo T, DePianto DJ, Salem O, Heigl T, Diamant N, et al. A cell atlas foundation model for scalable search of similar human cells. Nature. 2025;638(8052):1085-94.

7.	Hao Y, Stuart T, Kowalski MH, Choudhary S, Hoffman P, Hartman A, et al. Dictionary learning for integrative, multimodal and scalable single-cell analysis. Nature Biotechnology. 2024;42(2):293-304.
